# Cerebellar Granule Cells Develop Non-neuronal 3D Genome Architecture over the Lifespan

**DOI:** 10.1101/2023.02.25.530020

**Authors:** Longzhi Tan, Jenny Shi, Siavash Moghadami, Cydney P. Wright, Bibudha Parasar, Yunji Seo, Kristen Vallejo, Inma Cobos, Laramie Duncan, Ritchie Chen, Karl Deisseroth

## Abstract

The cerebellum contains most of the neurons in the human brain, and exhibits unique modes of development, malformation, and aging. For example, granule cells—the most abundant neuron type—develop unusually late and exhibit unique nuclear morphology. Here, by developing our high-resolution single-cell 3D genome assay Dip-C into population-scale (Pop-C) and virus-enriched (vDip-C) modes, we were able to resolve the first 3D genome structures of single cerebellar cells, create life-spanning 3D genome atlases for both human and mouse, and jointly measure transcriptome and chromatin accessibility during development. We found that while the transcriptome and chromatin accessibility of human granule cells exhibit a characteristic maturation pattern within the first year of postnatal life, 3D genome architecture gradually remodels throughout life into a non-neuronal state with ultra-long-range intra-chromosomal contacts and specific inter-chromosomal contacts. This 3D genome remodeling is conserved in mice, and robust to heterozygous deletion of chromatin remodeling disease-associated genes (*Chd8* or *Arid1b*). Together these results reveal unexpected and evolutionarily-conserved molecular processes underlying the unique development and aging of the mammalian cerebellum.

## Introduction

A broad goal of neuroscience is to achieve understanding of how different neural cell types mature even after birth along highly-distinct developmental trajectories (and, in a likely related question, how these cell types exhibit similarly diverse and specific patterns of degeneration during aging). Insight has recently emerged regarding the contributory molecular mechanism of diversity in genome architecture (cell-specific-specific folding of chromosomes in 3D to orchestrate gene transcription) (*1*). Although this process has been studied in both neurodevelopment and aging, genome architecture has never been systematically measured over the lifespan—limiting our understanding of molecular mechanisms that may be contributory to brain function and dysfunction.

Previously, using our high-resolution single-cell 3D genome assay (diploid chromosome conformation capture, or Dip-C) (*2–4*), we found that cells in the mouse forebrain (cerebral cortex and hippocampus) undergo profound, cell type–specific transformation in transcriptome and genome architecture during the first month of life—providing a structural foundation for functional maturation (*5*). However, it was unclear if this approach could generalize for comparison across brain regions, species (including human), and lifespan– a major set of goals that for quantitative and statistical accuracy would require new technology to profile many samples and donors in parallel in a single reaction, as well as new technology to isolate rare cell types that contribute to the differential cellular composition of neural circuits across region, species, and age.

Here we turn focus across the neuraxis, from forebrain to hindbrain– specifically to the cerebellum, which contains the majority (∼80%) of all neurons in the human brain. The cerebellum, an enormously powerful yet compact processing unit that has greatly expanded over the course of vertebrate and mammalian evolution (*6*), exhibits several unique biological characteristics. First, the cerebellum develops almost entirely after birth in mouse, and well into the first postnatal year in human. Second, it is the most consistently affected brain region in children with autism (*7, 8*), as well as a major locus of childhood cancer (medulloblastoma). Finally, the cerebellum undergoes substantial shrinkage with cell loss during aging (*9, 10*), but is one of the last regions to develop Alzheimer’s pathology. Understanding the molecular characteristics and genome dynamics of the cerebellum may provide invaluable insights into these unique features, as well as into the canonical functions of the cerebellum in motor control and cognition (including reward and sociability) (*11, 12*).

Prior correlative fluorescence and electron microscopy efforts have revealed unique nuclear morphological features of cultured mouse granule cells *in vitro* (*13*), but the 3D genome structures have never been solved. Recently, bulk chromosome conformation capture (3C/Hi-C) was performed on the whole adult mouse cerebellum (*14, 15*), and single-cell Hi-C on the adult human brain was reported in a preprint (*16*); however, a comprehensive, cross-species, single-cell 3D genome atlas of the developing and aging cerebellum—or any tissue for that matter—is still lacking. In addition, the relationships between 3D genome remodeling and transcriptome/chromatin accessibility changes during the early postnatal period have remained unknown in the cerebellum. Single-cell chromatin accessibility was profiled in the developing mouse and opossum cerebellum (*17*), and single-cell transcriptomes were profiled in the mouse cerebellum as well as in the developing human, mouse, and opossum cerebellum (*18*), but transcriptome and chromatin accessibility have never been simultaneously measured from the same cerebellar cells during development (nor has their relationship with 3D genome structure been analyzed).

Here we show that the mammalian cerebellum undergoes an extraordinary, life-long 3D genome transformation that is conserved between human and mouse and far greater in magnitude than any corresponding processes observed in the forebrain (*5*). We first created a multi-ome atlas of the developing human by simultaneously measuring transcriptome and chromatin accessibility within the same cell. In contrast with structural maturity by gross anatomy at birth, the newborn human cerebellum was found to harbor a continuous and broad spectrum of immature and mature granule cells in the first years of life. To understand the structural basis of this postnatal cerebellar maturation, we developed two new 3D genome technologies: population-scale Dip-C (Pop-C) to pool and profile many samples without batch effects, and virus-enriched Dip-C (vDip-C) to study rare cells (here, Purkinje cells that are critically important but present at <0.5% of the total population) without the need for transgenic mouse lines. Using Pop-C and vDip-C, we solved the initial 3D genome structures of single cerebellar cells, and created a high-resolution atlas for human and mouse spanning both development and aging. We found that although born with a 3D genome structure type similar to forebrain neurons (*5*), cerebellar granule cells unexpectedly transformed into a highly distinct structural type with specific inter-chromosomal contacts as well as ultra-long-range (10–100 Mb) intra-chromosomal contacts that had been thought to be exclusive to non-neuronal cells (e.g., microglia). In contrast to initial development, this 3D maturation proceeded continuously over lifespan like an aging clock, and was found to be highly robust in exhibiting resistance to functional perturbations including clinically-relevant heterozygous deletion of autism-implicated chromatin remodelers *Chd8* and *Arid1b*, and granule cell–specific deletion of *Chd4* (*15*). Together, this work has revealed not only an unprecedented genomic dimension of hindbrain maturation, but also the initial example of 3D genome atlasing across the lifespan, enabling discovery of lifelong genome rewiring as a new molecular hallmark of aging.

## Results

### A 3D genome atlas of the developing and aging cerebellum

In mouse, cerebellar granule cells (the vast majority of cerebellar neurons) are generated between postnatal days (P) 0–21; during this period, granule cell progenitors divide, mature, and migrate from the external granular layer (EGL) to the internal granular layer (IGL) of the cerebellar cortex, expanding >100-fold in number. In human beings, granule cells are generated between the third trimester of pregnancy and the age of ∼1 year after birth, in stark contrast with the cerebral cortex where neurons are generated well before birth (*5*); the cerebellum also exhibits a uniquely slow epigenetic aging clock based on bulk DNA methylation assays (*19*). To explore the genomic underpinnings of this unique timeline of cerebellar development and aging, we created the initial 3D genome atlas extending across the human and mouse lifespan, alongside a combined transcriptome and chromatin-accessibility atlas focused on human postnatal development (**Fig. 1A**).

**Fig. 1.**
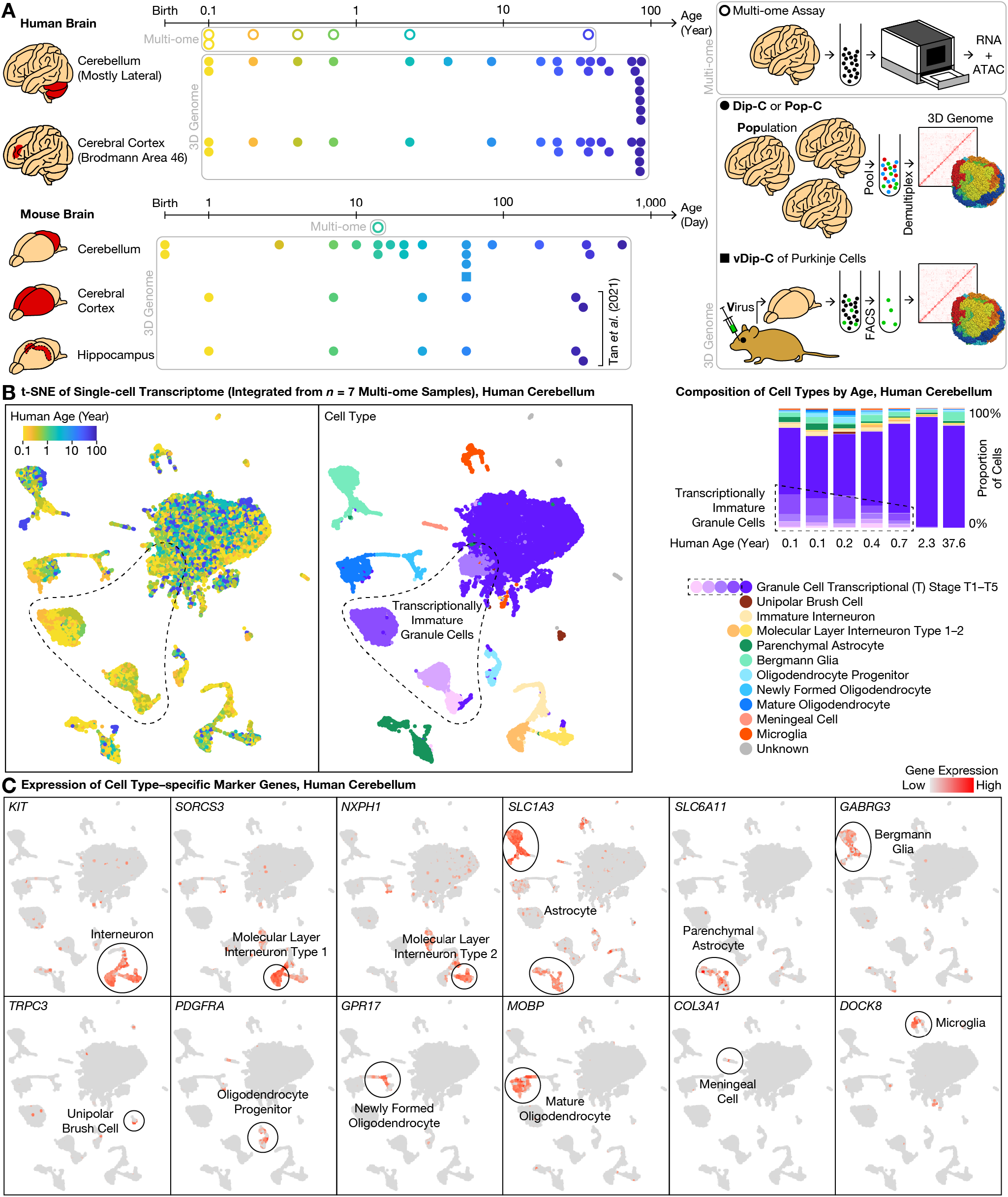
Single-cell 3D genome atlas across lifespan for human and mouse cerebellum with single-cell multi-ome atlas of postnatal cerebellar development. **(A)** Schematic of overall study design. To understand the genomic underpinning for cerebellar development and aging, we created a 3D genome atlas (filled dots and squares) of 13,161 cells from the cerebellum and cerebral cortex (and additionally, hippocampus in mouse) across the human (0.1–86 years) and mouse (0–2 years) lifespan, using our diploid chromosome conformation capture (Dip-C) (*2*), population-scale Dip- C (Pop-C), and virus-enriched Dip-C (vDip-C) methods (lower right). We additionally created a multi-ome (simultaneous transcriptome and chromatin accessibility) atlas (circles) of 63,768 cells from the developing cerebellum (human: 0.1–2.3 years and 1 adult; mouse: postnatal day (P) 14) (upper right; Methods). Data were compared to mouse cerebral cortex and hippocampus 3D genome data from previous work (*5*). **(B)** Integrative transcriptome analysis of the 7 human multi- ome samples revealed transcriptionally immature granule cells (dashed outlines) in the newborn cerebellum. We visualized the transcriptome portion of our human multi-ome atlas with t- distributed stochastic neighbor embedding (t-SNE) after cross-sample integration (*21*) (left: each dot represents a single cell). Granule cells existed in both mature form, which we termed transcriptional (T) stage 5 (T5; darkest purple), and a variety of immature forms, which we termed T1–T4 (dashed outlines; lighter shades of purple). Transcriptionally immature granule cells were abundant (14–34%) in the first postnatal year, but vanished (<1%) in the 2.3- and 37.6-year-old donors (right). **(C)** Representative gene expression profiles of cell type–specific marker genes (see **Fig. 2A** for granule cells).

We first simultaneously profiled transcriptome and chromatin accessibility during postnatal development of the human cerebellum, sequencing 63,768 cells from 7 donors (6 between the ages of 0.1–2.3, and 1 adult) and detecting a median of 645–1,617 genes (944–3,845 unique molecular identifiers (UMIs)) and 12–34 k assay for transposase-accessible chromatin (ATAC) fragments per cell from each donor (**Table S1, Table S2**). To compare between species, we additionally profiled a critical age in mouse—P14, where a substantial number of cells are present in both the EGL and the IGL—sequencing 7,182 cells and detecting a median of 618 genes (944 UMIs) and 22 k ATAC fragments per cell. Compared to existing transcriptome-only (*18*) or chromatin accessibility–only (*17*) data, this multi-ome dataset opens up new opportunities for cross-modal analysis during this highly dynamic window of cerebellar development.

We then comprehensively profiled 3D genome architecture across the human and mouse lifespan by developing 2 new 3D genome technologies (Pop-C and vDip-C; see below)—sequencing a total of 11,207 cells (**Fig. 1A**). In human, we sequenced 5,202 cells from 24 donors—spanning the ages of 0.1–86—and obtained a median of 608,000 chromatin contacts per cell. Among these cells, 3,580 came from the cerebellum (chiefly lateral; vermis if lateral not available), and 1,622 from the cerebral cortex (Brodmann area (BA) 46 of the dorsolateral prefrontal cortex (DLPFC)) of the same donors (**Table S2, Table S3, Table S4**). In mouse, we sequenced 6,005 cells from cerebellum—spanning the ages P0–P637 (birth to ∼21 months)—obtaining a median of 496,000 contacts per cell, and compared with our prior dataset of 1,075 and 879 cells from mouse cerebral cortex and hippocampus, respectively (*5*). With a combined cell number of 13,161, this 3D genome dataset represents the first life-spanning architectural atlas of any organ.

### Transcriptionally immature granule cells in the newborn (<1 year) human cerebellum

Validating cellular composition of the multi-ome atlas, this dataset accurately recapitulated known cerebellar cell types and marker genes. We integrated the transcriptome dataset from all 7 human donors with linked inference of genomic experimental relationships (LIGER) (*18, 20, 21*) (**Fig. 1B**); modifying LIGER parameters did not affect conclusions (**Fig. S1**). As expected, granule cells were the predominant cell type at all ages examined, representing a median of 83% (range: 76– 92%) of all cells. The next most abundant cell type, astrocytes (median: 7%; range: 3–11%), were composed of the cerebellum-specific *SLC1A3*-high/*AQP4*-low Bergmann glia (4%) (*20*) and the typical *SLC6A11*+/*SLC1A3*-low/*AQP4*-high parenchymal astrocytes (2%) (*6*)(*17, 18*) (**Fig. 1C**). We further identified *GABRG3*, a gamma-aminobutyric acid (GABA) receptor subunit, as a more specific marker for Bergmann glia. Less-abundant cell types included interneurons (4%), oligodendrocytes (3%), microglia (1%), and the relatively cerebellum-specific unipolar brush cells (UBCs) (0.3%). Note that due to sample size limitations, the cerebellum-specific Purkinje cells— critically important sole output neurons of the cerebellar cortex—were too rare to be reliably identified.

We focused initially on the granule cells—not only the most abundant cell type of the cerebellum, but also by cell number the dominant neuron type (∼80%) across the entire human brain, and due to their delayed generation critical for understanding functional maturation of the brain. Unlike the case of mouse brain wherein granule cells are almost entirely in the EGL at birth, the human cerebellum at birth has already attained relative maturity by gross anatomical measures, with IGL neurons outnumbering EGL neurons. However, it remains unknown when and how human granule cells mature at the molecular and genomic level after birth. We found that despite relative anatomical maturity of the human cerebellum at birth, granule cells were clearly subdivided into various maturation stages by transcriptomic measures (**Fig. 1B**). In our integrated transcriptome data, granule cells existed in both a mature form (which we term transcriptional (T) stage T5) and a variety of immature forms (stages T1–T4). The mature stage T5 was the predominant form (99– 100% of all granule cells) in the 2.3- and 37.6-year-old donors. In younger donors, however, the immature stages T1–T4 made up a substantial fraction of granule cells: 32–34% in the 0.1- and 0.2-year-olds, 23% in the 0.4-year-old, and 14% in the 0.7-year-old (**Fig. 1B**), revealing a surprising abundance of transcriptionally immature granule cells in the newborn human cerebellum.

The 5 transcriptional stages of granule cells expressed partially overlapping sets of genes (**Fig. 2A, Table S5**). Stage T1 was enriched for ribosomal subunits (FDR = 8 × 10^−67^; e.g., *RPS24/11/27* and *RPL31/39/32*) as well as axon guidance-related genes (FDR = 1 × 10^−4^; e.g., *BOC* and *LAMA2*), and was most specifically marked by *FOXP2*, a transcription factor (TF) implicated in speech development. Stage T2 was enriched for nervous system development-related genes (FDR = 3 × 10^−6^; e.g., *NFIB* and *UNC5C*) including additional genes related to morphogenesis of projections (FDR = 3 × 10^−6^; e.g., *ROBO2* and *MYO16*). Stage T3 was also enriched for nervous system development (FDR = 6 × 10^−12^; e.g., *ERBB4*) and projection morphogenesis (FDR = 1 × 10^−9^; e.g., *SEMA6D*), and was most specifically marked by *GRIA2*, a glutamate ionotropic receptor subunit implicated in multiple mental disorders. Stage T4 was enriched for cell adhesion (FDR = 3 × 10^−14^; e.g., *CNTNAP2/5* and *CNTN5*) and additional genes related to nervous system development (FDR = 2 × 10^−9^; e.g., *CHRM3* and *GPC6*). Finally, the mature stage T5 was enriched for synaptic signaling genes (FDR = 5 × 10^−16^; e.g., *CADPS2* and *SNAP25*) and regulation of membrane potential (FDR = 3 × 10^−11^; e.g., *RIMS1* and *SCN2A*), and was most specifically marked by *RBFOX1*, an RNA-binding protein implicated in many mental disorders. An alternative analysis— identifying 6 correlated gene modules from immature granule cell–enriched LIGER factors— confirmed and validated these genes and pathway enrichments (**Fig. S2, Table S5**). These stage-specific genes together help define the dynamics of complex transcriptional programs of postnatal cerebellar development that may be relevant to health and disease.

**Fig. 2.**
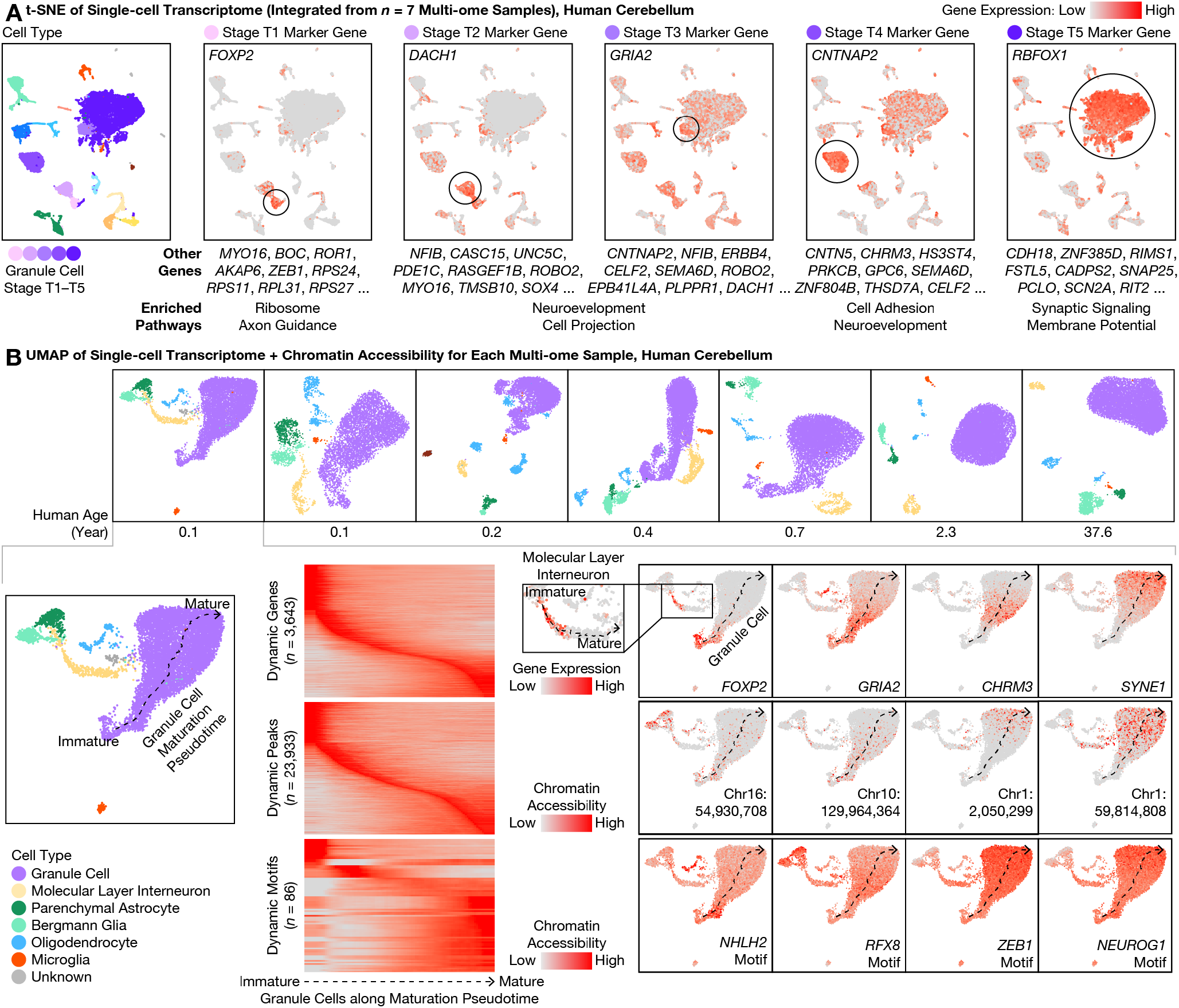
Simultaneous transcriptome and chromatin accessibility profiling revealed continuous maturation of cerebellar granule cells over the first postnatal year. **(A)** From our multi-ome atlas of the developing human cerebellum (**Fig. 1B**), we identified marker genes with expression enriched in each of the 5 transcriptional (T) stages (T1–T5; shades of purple) of granule cell maturation. Marker genes were ranked by specificity (area under the receiver operating characteristic (AUROC)). Expression of the top marker gene for each T stage was visualized on the integrated transcriptome t-SNE plot, while the top 2–10 genes were listed below. Enriched pathways (gene ontology (GO) terms) were summarized for the top 100 genes at each T stage. Note that overlap between marker genes at adjacent T stages suggested a continuous spectrum of transcriptional maturation. **(B)** We individually visualized each multi-ome sample with joint uniform manifold approximation and projection (UMAP) of transcriptome and chromatin accessibility (*22*), colored by cell types jointly defined by transcriptome and chromatin accessibility (top row). In each newborn sample (bottom: 0.1-year-old donor shown as example), granule cells (purple) exhibited a continuous maturation pseudotime (dashed arrows), suggesting gradual changes in both transcriptome and chromatin accessibility rather than discrete jumps. We identified dynamically expressed genes, dynamically accessible chromatin regions (peaks), and dynamically accessible transcription factor binding site (TFBS) motifs during granule cell maturation. Inset showed maturation pseudotime of molecular layer interneurons (MLIs).

### A continuum of granule cells and interneurons with maturing transcriptome and chromatin accessibility

For each donor, we next jointly analyzed the transcriptome and chromatin accessibility of each cell using analysis of regulatory chromatin in R (ArchR) (*22*) (**Fig. 2B**). Despite discrete appearances in LIGER’s cross-sample visualization (**Fig. 1A**), the 5 transcriptional stages T1–T5 of granule cell maturation formed a continuous developmental pseudotime for each donor below the age of 1—regardless of whether transcriptome and chromatin accessibility were analyzed alone or jointly (**Fig. S3, Fig. S4A**). In addition, ArchR identified genes, accessible chromatin regions (peaks),and TF binding site (TFBS) sequence motifs (motifs) that exhibited dynamic activity along the maturation pseudotime. Dynamically expressed genes included many genes from the above LIGER analysis, which turned on (and off for immature-only genes) at distinct pseudotimes. Dynamically accessible motifs included ASCL1/2 and NHLH1/2 motifs at early pseudotimes, KLF11/14 and RFX2/3/4/8 motifs at intermediate pseudotimes, and NEUROG1/2/3, NEUROD4/6, ZEB1, MEF2A/B/C/D, NFIA/B/X motifs at late pseudotimes. The human cerebellum in the first postnatal year was thus found to include a complex mixture of granule cells with continuously evolving transcriptomic and epigenomic states.

We validated aspects of this coexisting continuum of transcriptionally maturing granule cells in human cerebellum by re-analyzing transcriptome-only pre-printed data (*18*). Transcriptional cell types were concordant between the datasets; based on marker gene expression, our granule cell stages T1 (*BOC*+), T2 (*DDAH2*+), T4 (*CHRM3*+), and T5 (*RBFOX1*+) corresponded to granule cell progenitors (GCP), differentiating granule cells 1 (GC_diff_1), differentiating granule cells 2 (GC_diff_2), and defined granule cells (GC_defined) in (*18*), respectively, while our stage T3 (*ERBB4*+) corresponded to both GC_diff_1 and GC_diff_2. When each donor was re-analyzed individually, maturation stages displayed comparable overlap in marker gene expression in the two datasets (**Fig. S4B**), and immature granule cells vanished around the same ages in both studies—between 0.7–2.3 in our data and 0.8–2.8 in (*18*). Our dataset was therefore consistent with recently preprinted data, while further providing the epigenome dimension of information from the very same cells.

Our multi-omic observations from human brains were conserved in mouse. Indeed, a continuum of maturing granule cells was also observed in our mouse multi-ome sample at P14, with marker gene expression similar to our data from human cerebellum (**Fig. S5**), and to our re-analysis of preprinted transcriptome-only data (*18*). Unexpectedly, we noticed that one of the 5 granule cell sub-types (Granule_5) in a recent transcriptome atlas of the adult mouse cerebellum could in fact correspond to our immature granule cell stage T4 (*CHRM3*+), particularly given the relatively young (P60) age of the animals in that study (*20*). Interestingly, we observed a similar continuum of maturation in the second most abundant neuron type of the cerebellum—molecular layer interneurons (MLIs; median: 4%; range: 2–7%); in a pattern similar to that of granule cells, an immature MLI type was abundant at birth and vanished over age, comprising 64–71% of all MLIs in the 0.1- and 0.2-year-olds, 44% in the 0.4-year-old, 25% in the 0.7-year-old, 5% in the 2.3-year-old, and <1% in the adult (**Fig. 1B**). In addition, immature MLIs and immature granule cells shared expression of many genes, such as *FOXP2* (**Fig. 2B**). During postnatal development, we found MLIs to mature into the 2 distinct sub-types previously described in mouse (*18, 20*)—the gap junction-forming *PTPRK*+/*SORCS3*-high type 1, and the *NXPH1*+ type 2. We performed pseudotime analysis on MLIs and identified overlap with granule cells in dynamic genes, peaks, and motifs (**Fig. S6** and **Fig. 2B**), although a larger sample size is needed to compare rigorously.

### 3D genome profiling of diverse populations and rare cells with Pop-C and vDip-C

To understand the structural basis behind this postnatal transcriptome and epigenome transformation, we next focused on characterizing cerebellar 3D genome architecture across the human and mouse lifespan. Cerebellar cells exhibit unique genome morphology, beginning with nuclear dimension; for example, granule cell nuclei are among the smallest in the brain (∼6 mm diameter in rat), while Purkinje cells have relatively large nuclei (∼12 mm diameter in rat) (*23*). Moreover, during differentiation, cultured mouse granule cell progenitors rapidly reduce nuclear (specifically euchromatin) volume and spatially redistribute histone H3.3 (*13*). However, 3D genome structures of cerebellar cells have not been resolved and little is known about lifetime- spanning dynamics *in vivo*.

We developed 2 new single-cell sequencing technologies to enable high-throughput, reproducible creation of a comprehensive cerebellar 3D genome atlas. First, population-scale Dip-C (Pop-C) leveraged the whole-genome sequencing capability of our Dip-C method (*2*) to pool a large number of samples/donors in a single reaction, and computationally demultiplexed (*24*) individual cells based on their natural variations in linear DNA sequences (**Fig. 1A, Fig. 3A**). By pooling tissues before homogenization, Pop-C eliminated nearly all batch effects that have previously plagued single-cell studies, and at the same time substantially reduced time, labor, and reagent cost. Compared to pooled transcriptome profiling (*25*) from which we took inspiration, Pop-C was even better suited for demultiplexing because of its high genomic coverage. We validated the performance of Pop-C by computationally pooling known samples as gold standards.

**Fig. 3.**
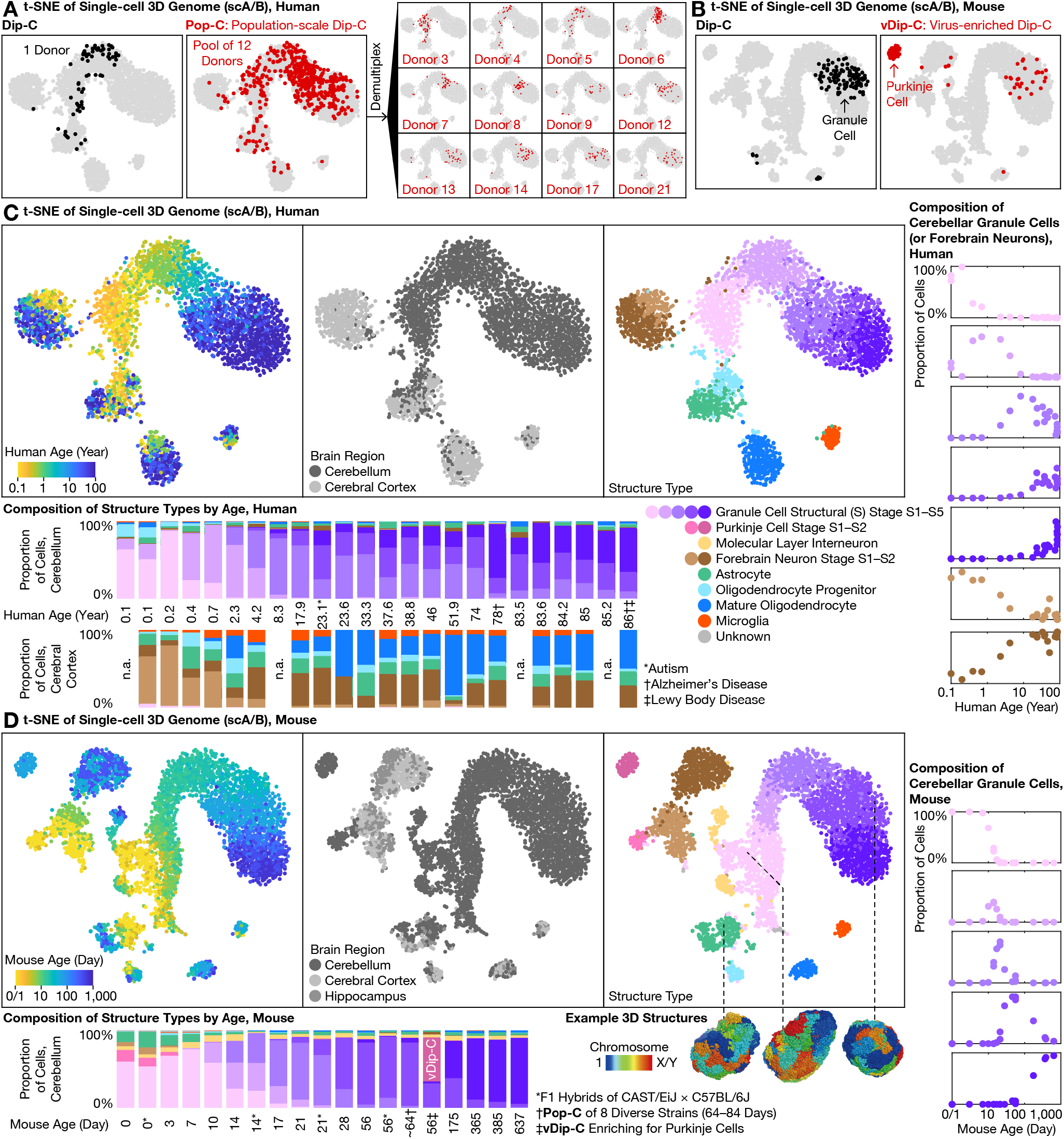
High-throughput, high-precision 3D genome profiling uncovered progressive lifelong genome remodeling in the human and mouse cerebellum. **(A)** Traditional 3C/Hi-C methods such as Dip-C measure one sample at a time (left); in contrast, the new population-scale Dip-C (Pop-C) method simultaneously profiles many samples at once, by pooling samples and computationally demultiplexing single cells based on linear DNA sequences (genotypes), thereby eliminating batch effects and substantially reducing labor and cost (right). This approach was found to be highly reproducible; note consistency between the same sample (Donor 4) assayed both individually (left) and by Pop-C (right), visualized with t-SNE of the single-cell 3D genome (based on single-cell chromatin A/B compartment values; scA/B) (*5*); each dot represents a single cell). **(B)** Rare cell types, such as adult Purkinje cells (<0.5% in mouse), were often missed by conventional Dip-C (left); in contrast, virus-enriched Dip-C (vDip-C) enabled efficient isolation of adult Purkinje cells with minimally invasive, fixation-robust fluorescent labeling in wild-type mouse tissue (right). **(C**–**D)** With Pop-C and vDip-C, we created a high-resolution, cross-species 3D genome atlas for the developing and aging cerebellum (with cerebral cortex as counterpoint), and resolved the first 3D genome structures of single cerebellar cells (bottom (D)). In both species (human (C); mouse (D)), all samples were visualized together with t-SNE of scA/B; 3D genome structure types were identified with hierarchical clustering of scA/B (top). Cerebellar granule cells (shades of purple) exhibited by far the most dramatic structural transformation—born with an immature structure type, which we termed structural (S) stage S1 (lightest purple), that closely resembled forebrain neurons (shades of brown), and evolving progressively into new structure types, which we termed stages S2–S5 (darker shades of purple), that drastically differed from all other neurons as the cerebellum developed and aged. Abundances of the various S stages peaked around ages 0.2, 1, 10, 30, and 80 yr in human, and around P3, P14, P21, P56 (∼2 months), and P365 (∼12 months) in mouse (bottom, right).

Pop-C reliably pooled and demultiplexed a variety of human and mouse samples. In the simplest case, many of our mouse samples were a pool of 1 male and 1 female of the same age, which we demultiplexed based on the ratio of read (or contact) densities between X chromosome and autosomes (termed X:A). In a more complex case, we pooled 1 young adult (P56–84) male mouse each from the 8 founder strains of the JAX Diversity Outbred (DO) collection (*26*)—A/J, C57BL/6J, 129S1/SvImJ, NOD/ShiLtJ, NZO/HlLtJ, CAST/EiJ, PWK/PhJ, and WSB/EiJ—which we demultiplexed based on their known single-nucleotide polymorphisms (SNPs) from the Mouse Genomes Project (**Table S4**). In the most challenging case, we pooled 3–13 unrelated human individuals and demultiplexed them with souporcell (*24*) based on known common SNPs in human populations, without prior knowledge about donor genotypes (**Fig. 3A**). Pop-C was thereby shown to provide a convenient, robust, and rigorous method for profiling single-cell 3D genome at unprecedented scales (**Fig. S7**).

Virus-enriched Dip-C (vDip-C), on the other hand, enabled genomic profiling of rare cell populations without the need for transgenic mouse lines (**Fig. 1A, Fig. 3B**). Testing a previous methodology involved isolating 3 cortical neuron types for chromatin accessibility and DNA methylation by genetically targeting a fluorescent protein to the nuclear membrane (*27*); however, this required a transgenic mouse line (and in that case, crossing with another line for cell type– specific Cre expression). Moreover, this approach is vulnerable to loss of fluorescence after formaldehyde fixation that would adversely affect many epigenome assays such as Dip-C. Here we developed vDip-C—a compact, single-component viral vector containing a cell type–specific promoter, an ultra-bright, fixation-resistant, monomeric fluorescent protein mGreenLantern (*28*), and a short (∼250 bp) nuclear membrane localization sequence (KASH domain of the mouse *Syne2* gene) (*29*) (**Fig. S8**). We found that this vector could be conveniently and minimally invasively administered to wild-type mice (e.g., C57BL/6J and MOLF/EiJ (*30*)) via retro-orbital injection of a blood-brain barrier (BBB)–crossing adeno-associated virus (AAV) PHP.eB serotype (*31, 32*).

We used vDip-C to solve the first 3D genome structures of single Purkinje cells (**Fig. 3B**). Although Purkinje cells are relatively abundant at mouse birth (P0), they quickly become outnumbered by the rapidly dividing granule cells (**Fig. 3D**). To isolate this rare (<0.5%) cell type from the adult mouse cerebellum, we constructed a vDip-C vector with a Purkinje cell–specific promoter (from the mouse *Pcp2* (also known as the L7) gene (*33*)), administered to wild-type mice—either strain C57BL/6J or the filial 1 (F1) hybrid of C57BL/6J and MOLF/EiJ (*30*) for 3D genome reconstruction—and isolated nuclei by fluorescence-activated cell sorting (FACS) (**Fig. S8**). As more cell type–specific viral promoters/enhancers are now being developed and discovered, vDip-C will increasingly provide a readily-applicable and general tool for epigenome profiling of many brain cell types.

### Life-long 3D genome transformation of granule cells

With Pop-C and vDip-C, we created a high-resolution, cross-species single-cell 3D genome atlas for the developing and aging cerebellum—using the cerebral cortex for comparison—and resolved 3D genome structures for a subset of cells (from F1 hybrid mice) (**Fig. 1A, Fig. 3C, Fig. 3D**). Similar to our previous studies in the mouse forebrain (*5*) and in other systems (*2, 3*), this new single-cell chromatin A/B compartment (scA/B) analysis revealed 3D genome structure types corresponding to diverse cerebellar cell types—including granule cells, astrocytes, oligodendrocytes, and microglia in both species, as well as MLIs and Purkinje cells in mouse. These 3D genome measurements were highly robust, in that replicates (e.g., multiple tissue dissections, multiple Dip-C/Pop-C batches) of the same donors yielded consistent scA/B patterns (**Fig. 3A, Fig. S7**). Note that 3 of our 24 human donors were diagnosed with autism, Alzheimer’s disease, and/or Lewy body disease; excluding these donors did not affect our conclusions.

Among all cell types examined, cerebellar granule cells exhibited by far the most dramatic structural transformation. We previously reported extensive 3D genome rewiring in the mouse forebrain (cerebral cortex and hippocampus) after birth (*5*) (brown cells in **Fig. 3D**); however, the magnitude of forebrain changes was dwarfed by changes in cerebellar granule cells (purple cells). In particular, granule cells of both species were born with an immature structure type, which we termed structural (S) stage S1, that most closely resembled that of forebrain neurons (**Fig. 3C, Fig. 3D**). As the cerebellum developed and aged, granule cells continuously and progressively evolved into a new structure type—going through stages S2–S5—that drastically differed from all other neurons. This 3D genome transformation represented the primary source of scA/B variations (i.e., the first principal component (PC)) in our atlas for both species, and therefore could be prominently visualized regardless of the dimension reduction method—principal component analysis (PCA), t-distributed stochastic neighbor embedding (t-SNE), or uniform manifold approximation and projection (UMAP) (**Fig. S9**).

Life-long 3D genome changes in granule cells followed a continuously progressing trajectory across the lifepsan. To facilitate analysis, we clustered granule cells into 5 structural (S) stages S1– S5 based on scA/B. In human, abundances of S1–S5 peaked around the ages of 0.2, 1, 10, 30, and 80, respectively, although considerable between-donor variability was observed (**Fig. 3C**). In mouse, S1–S5 peaked around P3, P14, P21, P56 (∼2 months), and P365 (∼12 months) (**Fig. 3D**); note that although not directly captured by our 5-stage analysis, mouse 3D genome structure continued to mature between P365/385 (∼12 months) and P637 (∼21 months) (**Fig. S10**). These data therefore reveal the initial example of a 3D genome aging clock, revealed by uncovering a surprisingly large architectural transformation in a mostly post-mitotic cell type.

### Ultra-long-range intra-chromosomal contacts and specific inter-chromosomal contacts in granule cells

The most prominent architectural changes in granule cells we observed were the emergence of ultra-long-range (10–100 Mb) intra-chromosomal contacts that had previously been thought exclusive to non-neuronal cells (**Fig. 4A**). In both species, the fraction of intra-chromosomal contacts in granule cells that were ≥10 Mb steadily increased across structural stages S1 and S5— from (19 ± 4)% to (33 ± 3)% (mean ± s.d.) in human, and from (19 ± 6)% to (34 ± 2)% in mouse. Such prevalence of ultra-long-range contacts was found to be in sharp contrast to that of forebrain neurons (which were observed to be only changing from (15 ± 4)% to (16 ± 5)% during human development, and from (11 ± 4)% to (13 ± 3)% in mouse), and of Purkinje cells (from (9 ± 1)% to (10 ± 2)% in mouse).

**Fig. 4.**
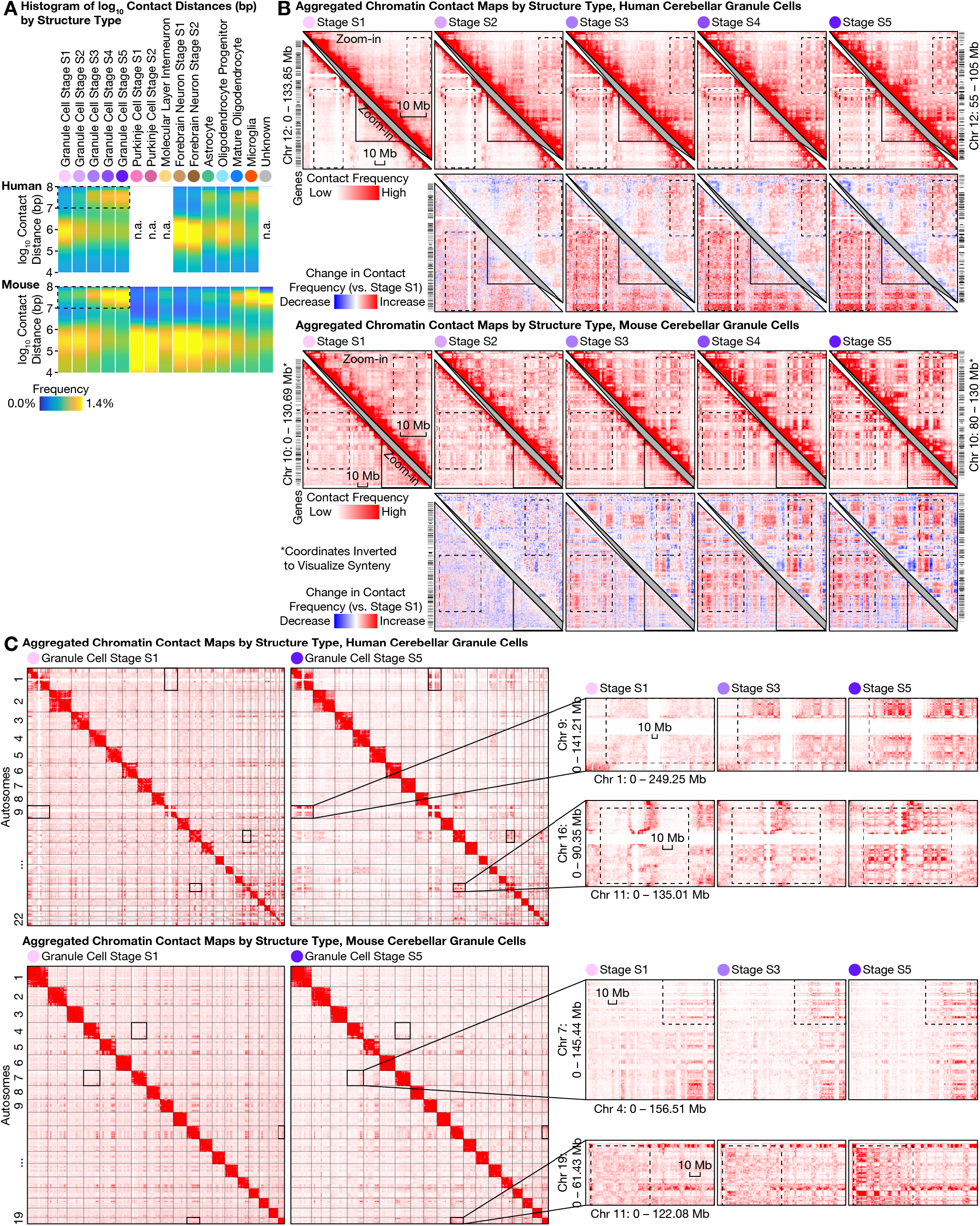
Large-scale 3D genome remodeling of cerebellar granule cells formed ultra-long-range (10–100 Mb) intra-chromosomal contacts and specific inter-chromosomal contacts during development and aging. **(A)** The most prominent architectural changes in granule cells (first 5 columns) were emergence of ultra-long-range (10–100 Mb) intra-chromosomal contacts (dashed boxes) thought to be exclusive to non-neuronal cells (*16, 34*) such as microglia (second to last column). Distribution of genomic distances of chromatin contacts (in base pairs (bp); on logarithmic scale) quantified by histogram for each 3D genome structure type (top) in human (middle) and mouse (bottom). **(B)** Emergent ultra-long-range contacts in granule cells formed prominent checkerboard patterns on contact maps—suggesting strong phase separation between the newly formed chromatin A/B compartments; this effect was generally stronger in the gene-poor, heterochromatic compartment B. For each species, an aggregated contact map of each structural (S) stage (S1–S5) of granule cell maturation was shown for an example chromosome (lower left triangles) and for an example zoomed-in (50-Mb) genomic region (upper right triangles). Contact maps (matrices of contact frequencies) were visualized with Juicebox (*47*), both as absolute values (first and third rows) and as relative changes compared to stage S1 (second and fourth rows). Zoomed-in regions are homologous between human and mouse; mouse coordinates were inverted for synteny. Dashed boxes highlight prominent changes during granule cell maturation. Bin size: 250 kb. **(C)** Granule cells formed specific inter-chromosomal contacts during development and aging; note increasing interactions between certain chromosomes—most prominently within a multi-chromosome hub of Chr 1/9/11/14/15/16/17/21/22, and between chromosome pairs such as Chr 2/9, Chr 4/14, Chr 8/11, Chr 13/20 in human. In each species, aggregated contact maps are shown for 2 example chromosome pairs. Dashed boxes highlight prominent changes during granule cell maturation. Bin sizes: 6 Mb (human genome-wide); 5 Mb (mouse genome-wide); 500 kb (zoom-in).

This progression in granule cells ended in a state that curiously more closely resembled non-neuronal cells such as microglia ((34 ± 3)% in both species) and mature oligodendrocytes ((29 ± 4)% in human, (27 ± 5)% in mouse). These ultra-long-range contacts formed prominent checkerboard patterns on contact maps—suggesting phase separation between newly formed chromatin A/B compartments—and were generally stronger in the gene-poor, heterochromatic compartment B in granule cells (**Fig. 4B**). Further heightening the novelty of this finding, two preprints reinforced the dichotomy between the compartment-dominant (i.e., longer-range) 3D genome in non-neuronal cells and the domain-dominant (i.e., shorter-range) 3D genome in neurons and their progenitors (radial glia) across the human body (*16, 34*). Our results therefore provide, in cerebellar granule cells, a striking counter-example of neurons converting from a neuronal structure type to one of the strongest non-neuronal structure types via postnatal development and aging.

Unexpectedly, granule cells’ redistribution of intra-chromosomal contacts was accompanied by the formation of highly specific inter-chromosomal contacts. We found increasing interactions among certain human chromosomes—most prominently within a multi-chromosome (Chr) hub of 1, 9, 11, 14, 15, 16, 17, 21, and 22, and between chromosome pairs such as Chr 2/9, Chr 4/14, Chr 8/11, Chr 13/20; meanwhile, Chr 12 weakened its interactions with several chromosomes in the hub (e.g., with Chr 1, 9, 11, 16) (**Fig. 4C, Fig. S11**). Similar to the ultra-long-range intra- chromosomal contacts (**Fig. 4B**), these emergent, granule cell–specific inter-chromosomal contacts were generally stronger in the heterochromatic regions—suggesting a common mechanistic origin. Finally, we found these inter-chromosomal contacts conserved in mouse, despite extensive chromosomal re-arrangements during evolution (**Fig. 4C**); for example, the centromeric portion of mouse Chr 7 gained interactions with portions of Chr 4, 5, 11, 17, 19. Together, these results demonstrated another example of conserved, specific inter-chromosomal interactions aring in neurodevelopment, beyond prior discoveries in nasal tissue (*3, 35–37*).

### Life-spanning scA/B changes associated with granule cell–specific marker genes

To understand the relationship between our observed transcriptional and architectural changes in granule cells, we examined scA/B dynamics of each genomic locus in detail. We have previously shown that scA/B generally correlates with cell type–specific gene expression in a variety of systems (the mouse eye, nose, cerebral cortex, and hippocampus), although discordance can be observed at the single-gene level and regarding temporal dynamics (*3, 5*). It remains unclear how scA/B interacts with gene expression during aging.

In granule cells, we found the predominant mode of scA/B changes to be progressive up- or down- regulation throughout postnatal development and aging. We calculated the mean scA/B of each 1- Mb genomic region (autosomes only) at each of the 5 structural stages S1–S5, and identified the top 20% dynamic regions based on between-stage variance. In both species, these ∼500 dynamic regions either gradually increased (becoming more euchromatic/compartment A–like) or decreased (more heterochromatic/B-like) in scA/B across the 5 stages—in a continuously progressing manner over age (**Fig. 5A**).

**Fig. 5.**
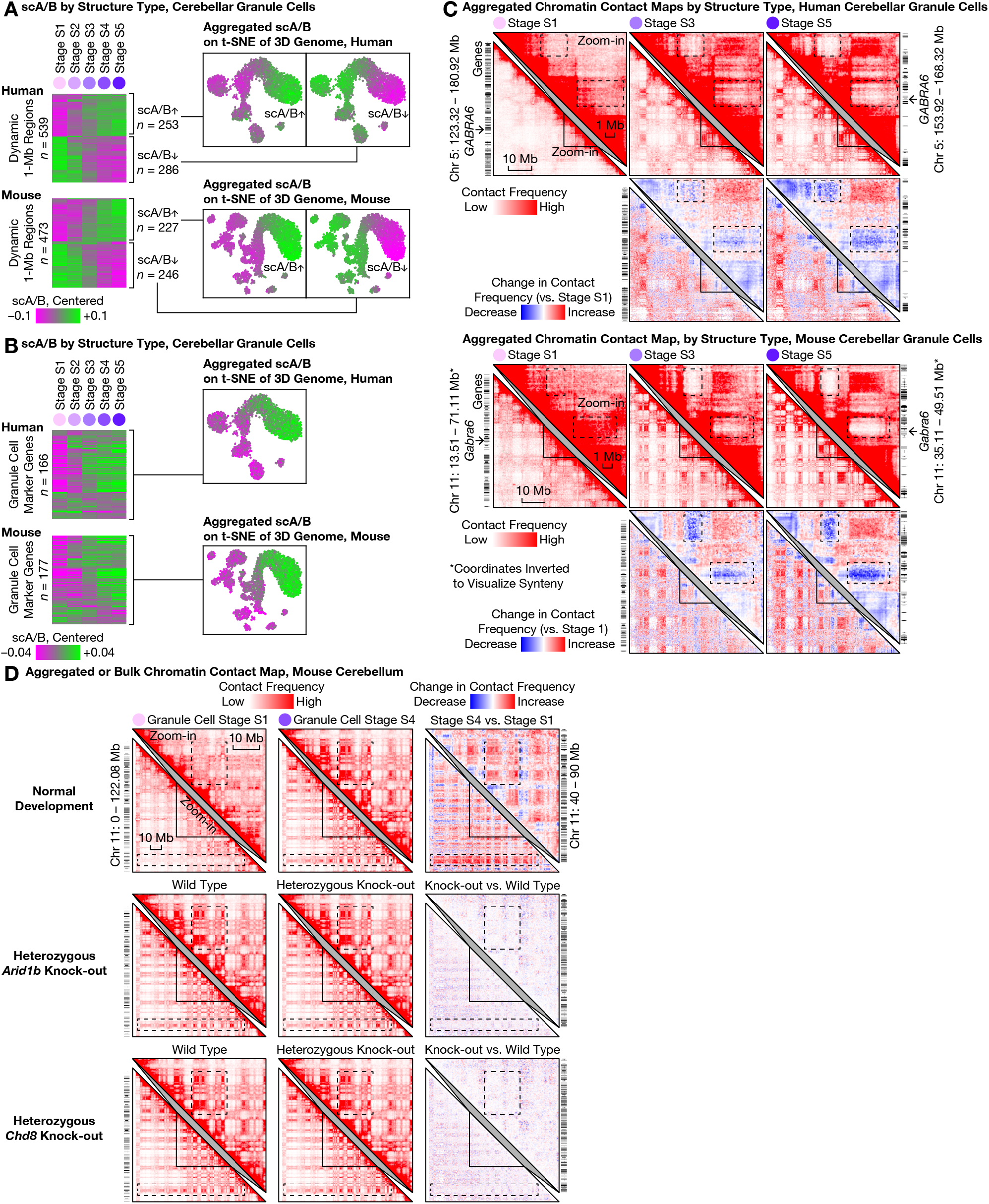
Continuous lifelong maturation of single-cell chromatin A/B compartment value (scA/B) was associated with cerebellar granule cell–specific marker genes and resistant to modulation of *Arid1b* or *Chd8*. **(A)** Relationship between transcriptional and architectural changes in granule cells. We calculated the mean scA/B of each 1-Mb genomic region at each structural (S) stage (S1–S5; columns) of granule cell maturation, and identified the top 20% dynamic regions (rows) based on between-stage variance. Dynamic regions (rows) are shown in a heatmap (left) ordered by hierarchical clustering of scA/B correlation, and clustered into 2 temporal modes: continuous up- or down-regulation across the lifespan. Each mode was additionally visualized with aggregated scA/B of all its regions on the 3D genome t-SNE plot (right). **(B)** Continuously-progressing scA/B changes correlated with expression of mature granule cell–specific genes—suggesting continued 3D genome rewiring well after initial transcriptional up-regulation. Mean scA/B of each 1-Mb genomic region harboring conserved, granule cell– specific marker genes (Supplementary Table 5 of (*18*)) at each S stage reveals that the majority of such regions continually increased scA/B across the lifespan (left). Shown: aggregated scA/B of all such regions on the 3D genome t-SNE plot (right). **(C)** The granule cell–specific marker gene *GABRA6* (arrows) steadily increased scA/B by losing contacts with two nearby gene-poor regions over the lifespan (dashed boxes; note the gene-poor regions formed strong contacts with each other over time) in both species. Contact maps were visualized with Juicebox (*47*), both as absolute values (first and third rows) and as relative changes compared to stage S1 (second and fourth rows). Zoom-in regions homologous between human and mouse; mouse coordinates inverted for synteny. Bin size: 250 kb. **(D)** Functional perturbation by disrupting chromatin remodelers. Bulk Dip-C on whole adult cerebellum (chiefly granule cells) of mice with clinically-relevant heterozygous deletion of autism-implicated genes *Arid1b* (middle) or *Chd8* (bottom) had little effect on 3D genome (top; stage S4 chosen to match ages). Contact maps visualized both as absolute values (left and middle) and as relative changes (right). Dashed boxes highlighted prominent changes during granule cell maturation. Bin size: 250 kb.

Interestingly, the continuously progressing scA/B changes were correlated with expression of mature granule cell–specific genes, suggesting continued 3D genome rewiring well after initial transcriptional up-regulation. We examined genomic regions that harbored conserved marker genes of mature granule cells from (*18*) (GC_defined, corresponding to our transcriptional stage T5 (**Fig. 1B**)); conclusions were robust to choice of marker gene set. Expression of these ∼200 marker genes began around birth, when transcriptionally mature granule cells (stage T5) emerged and replaced immature ones (T1–T4). However, we found that on average, the marker genomic loci gradually increased scA/B throughout life, i.e., from structural stages S1–S5 (**Fig. 5B**). For example, the mature granule cell–specific GABA receptor subunit *GABRA6* gradually lost contacts with 2 nearby gene-poor (heterochromatic) regions over the entire lifespan (**Fig. 5C**) and consequently, steadily increased scA/B from structural stages S1–S3 (∼10 years) in human, and from S1–S4 (∼P56) in mouse—persisting well after initial transcriptional up-regulation in both species (∼0.5 years and ∼P10, respectively (*18*)). Other example genes can be found in **Fig. S12**. In contrast, transcriptionally down-regulated genes, i.e., granule cell progenitor–specific marker genes (GCP in (*18*), or our transcriptional stage T1), on average exhibited relatively unchanged scA/B across structural stages S1–S5—consistent with previous observations in mouse nasal tissue (*3*). These results may contribute to molecular understanding of a curious temporal mismatch between transcriptional and structural changes, where increased transcription may continue to etch 3D genome structure long after initial gene activation as initially suggested by correlation between neuronal marker genes (activated before birth) and postnatal 3D rewiring in the mouse forebrain (*5*).

### Robust 3D genome maturation despite functional perturbations

To test robustness of this genome restructuring, we explored functional perturbation of granule cell chromatin remodeling, both in the setting of disrupted disease-implicated chromatin remodelers as well as via comprehensive re-analysis of published data (*14, 15*). *Arid1b* (*38*) and *Chd8* are two of the most frequently mutated genes in autism patients, encode chromatin remodelers, and are widely expressed in many cell types—including granule cells—of the developing and adult cerebellum (*18*). To test involvement in our observed architectural maturation, we performed bulk Dip-C on the whole adult cerebellum (P57–70, consisting predominantly of granule cells) of mice with clinically-relevant heterozygous deletion of *Arid1b* or *Chd8*. Despite relatively deep sequencing (313, 189, 219, and 231 million contacts for *Arid1b* heterozygote/control and *Chd8* heterozygote/control, respectively), little change in 3D genome organization was observed (**Fig. 5D**). We finally re-analyzed available bulk Hi-C data of perturbed cerebellum (**Fig. 5D, Fig. S13**). Conditional, homozygous deletion of *Chd4*—a chromatin remodeler implicated in intellectual disability and expressed in many cerebellar cell types—in mature granule cells (driven by *Gabra6-*Cre) caused moderate 3D changes across the mouse genome (*15*). Strikingly, however, these changes had little overlap with (and were of a much smaller magnitude than) our observed architectural maturation, revealing the 3D genome transformation to be highly robust in the setting of diverse chromatin perturbations.

## Discussion

Once born, most neurons in the central nervous system must last a lifetime, and adapt relevant gene expression programs to changing needs across the lifespan. However, we know surprisingly little about how the underlying genomic information is structurally organized over the lifespan, and specifically how this changing organization relates to gene transcription. Here we addressed this longstanding question in a biological system (the mammalian cerebellum) that is critically important in diverse adaptive and maladaptive brain states, under-studied with modern omics methods, and remarkable for its unique developmental trajectories and transformations.

Despite the increasingly-appreciated roles of the cerebellum in cognition, along with its well-characterized neural circuitry and physiology, recent large-scale neurogenomics efforts have focused on other brain regions such as the cerebral cortex, leaving the unique genomics of ∼80% of our brain’s neurons—the cerebellar granule cells—under-explored. By resolving these genome structures for the first time, including over the mammalian lifespan from birth to advanced age, we found unprecedented genome architecture: ultra-long-range intra-chromosomal contacts that blurred the genome-architectural boundary between neurons and non-neuronal cells, highly specific inter-chromosomal contacts reminiscent of those in nasal tissue (*3*), and continuous 3D remodeling over decades that may be stabilized by cell type–specific gene transcription. Along the way, we showed that mouse is an excellent animal model of this process, despite substantial differences from human beings in developmental timing and lifespan, by validating and extending the critical findings from mouse across the human lifespan.

The data presented here provide critical mechanistic insights into the fundamental principles of 3D genome reorganization. For example, previously in the mouse forebrain, we had discovered neuron-specific, large-scale inward movement of many genomic regions that is concurrent and inversely-correlated with neuron-specific non-CpG DNA methylation (*5*). Here we found that cerebellar granule cells also exhibit the same inward genome movement (**Fig. S14**), but since they curiously do not gain non-CpG methylation (according to a recent preprint(*16*), methylome changes may be dispensable for radial genome movement. Furthermore, our present results also highlight an intriguing juxtaposition of the uniquely non-neuronal 3D genome and non-neuronal methylome specific to granule cells; the lack of non-CpG methylation alone was insufficient to impose non-neuronal 3D genome architecture in the similarly un-methylated Purkinje cells. Finally, our results revealed robustness of the unusual granule cell genome architecture to strong functional perturbations such as clinically-relevant disruption of disease-implicated chromatin remodelers.

A potential function of this unique genome architecture might be to manage space and energy expenditure. Our brains are 80% cerebellar granule cells at least by neuron number–a fraction that has increased steadily in the lineage leading to modern humans, suggestive of an adaptive value to large numbers of granule cells. However, if each granule cell consumed the same volume and energy as a typical neuron in the cerebral cortex (e.g., a pyramidal cell), metabolic costs could become maladaptive. Consistent with this idea, granule cells are known to be relatively quiet by firing rate (∼0.1 Hz in mouse) (*39*), in contrast to the large and relatively active Purkinje cells (∼50 Hz in mouse) (*40*). Cerebellar granule cells might therefore have adopted a more compact non-typically neuronal form for both cell body and genome, and a more quiet state both physiologically and transcriptionally, in order to conserve space and energy. Considering these results alongside our previous work (*5*), it is worth noting that cerebellar and hippocampal granule cells adopt utterly different strategies of genome organization, despite sharing similar anatomy for dense packing (and hence similar nomenclature) in tissue. Indeed, per our 3D genome atlas, mouse hippocampal granule cells are more similar to other forebrain neurons than to cerebellar granule cells, for example in not developing ultra-long-range contacts. Small nuclear size might be the most powerful driver of non-neuronal genome architecture, since hippocampal granule cells have much larger nuclei (9–10 mm in human and in P70 rat) (*41, 42*) than cerebellar granule cells (5–6 mm in rat), although both cell types are similarly sparsely active (firing rate 0.1–0.2 Hz in mouse) (*43*). It remains to be determined how a third type of granule cells—olfactory bulb granule cells (which notably are inhibitory rather than excitatory)—organize and maintain their genome over the course of development and aging.

More broadly, this approach showcases how life-spanning 3D genome profiling of a complex, living tissue can provide unprecedented dimensions of information relevant to the molecular mechanisms of development and aging. This presence of a lifelong structural transformation may point the way to new therapeutic targets for interventions that could be specific to developmental and aging-related disorders. Wide application of the new 3D genome profiling technologies (Pop- C and vDip-C), along with their corresponding analysis pipelines, to many brain regions and tissues of the human body may contribute to solving longstanding challenges such as dissecting the genetic basis of inter-individual 3D genome variability, characterizing ultra-rare cell types, and revealing the full extent of the diversity and dynamics of 3D genome organization.

## Acknowledgments

We thank Arima Genomics for early access to the 3C kit, the National Institute of Mental Health (NIMH) Human Brain Collection Core (HBCC), the Stanford Alzheimer’s Disease Research Center (ADRC), and the National Institutes of Health (NIH) NeuroBioBank (NBB) for postmortem human samples, Feng Zhang (MIT) and Xin Jin (Scripps) for advice on *Chd8* mice and the KASH tag, Cemre Celen (UT Southwestern) for help on *Arid1b* mouse genotyping, Haynes Heaton (Auburn) for advice on souporcell, Ryan Corces (UCSF) for help with ArchR, Dong Xing (Peking), Jennifer Raymond (Stanford), Tom Clandinin (Stanford), Anne Brunet (Stanford), Linlin Fan (Stanford), Peter Wang (Stanford) for helpful discussion, students of the Stanford Immersive Neuroscience (SIN) bootcamp (Jacqueline Bendrick, Jerry Cheng, Cheyanne Lewis, Karen Malacon, Nick Manfred, Blake Zhou) for help on some Dip-C experiments, the Stanford PAN Facility, and the Stanford Shared FACS Facility. We thank the donors and their families, the Medical Examiners Offices in D.C., Richmond, and Northern Virginia for their contribution to the HBCC, the University of Maryland Brain and Tissue Bank (https://www.medschool.umaryland.edu/btbank/) and the Stanley Medical Research Institute Brain Research Tissue Repository (https://www.stanleyresearch.org/brain-research/) for contributing some tissues to HBCC.

## Funding

Burroughs Wellcome Fund (BWF) Career Award at the Scientific Interface (CASI) (LT)

Stanford University School of Medicine Dean’s Postdoctoral Fellowship (LT)

Stanford University Walter V. and Idun Berry Postdoctoral Fellowship (LT)

Stanford University Undergraduate Summer Research Fellowship in Chemistry (JS)

Stanford University Bio-X Undergraduate Summer Research Program (JS)

National Institute of Mental Health (NIMH) grant R01 MH123486 (LD)

National Institute of Mental Health (NIMH) grant R21 MH125358 (LD)

Stanford University Jaswa Innovator Award (LD)

National Institute of Neurological Disorders and Stroke (NINDS) grant K99 NS119784 (RC)

Gatsby Foundation grant (KD)

## Author contributions

Designed the experiments: LT, JS, IC, LD, RC, KD

Performed the experiments: LT, JS, BP, KV, RC

Analyzed the data: LT, JS, SM, CPW, BP, YS, KV, IC, LD, RC, KD

Wrote the manuscript: LT, KD

## Data and materials availability

Raw and processed data is available from the National Center for Biotechnology Information (NCBI) under the BioProject PRJNA933352 (https://www.ncbi.nlm.nih.gov/bioproject/?term=PRJNA933352). Code is available from GitHub (https://github.com/tanlongzhi/dip-c). The vDip-C vector will be available from Addgene.

