## Supplementary Materials for "Cerebellar Granule Cells Develop Non-neuronal 3D Genome Architecture over the Lifespan"

**The PDF file includes:**

Materials and Methods
Figs. S1 to S13
Tables S1 to S5

### Materials and Methods

Postmortem Human Samples

Fresh frozen, postmortem human brain samples were obtained from 3 tissue banks: the National Institute of Mental Health (NIMH) Human Brain Collection Core (HBCC), the Stanford Alzheimer's Disease Research Center (ADRC), and the National Institutes of Health (NIH) NeuroBioBank (NBB) (**Table S1**).

Animals

Animals were maintained in accordance with the U.S. National Research Council Guide for the Care and Use of Laboratory Animals. Animal protocols were approved by the Administrative Panel on Laboratory Animal Care (APLAC) at Stanford University. Animals (up to 5 per cage) were housed in an enriched environment on a 12 h dark/12 h light cycle. All animals had normal health/immune status, were not involved in previous procedures, and were drug-/test-naive.

The majority of mice were from the inbred C57BL/6J strain (JAX 000664). For 3D genome modeling, F1 hybrids were generated in house by crossing CAST/EiJ (JAX 000928) with C57BL/6J (JAX 000664) for regular Dip-C, or crossing MOLF/EiJ (JAX 000928; father) with C57BL/6J (JAX 000664; mother) for vDip-C. The 8 founder strains of the JAX DO collection were purchased from JAX: A/J (JAX 000646), C57BL/6J (JAX 000664), 129S1/SvImJ (JAX 002448), NOD/ShiLtJ (JAX 001976), NZO/HlLtJ (JAX 002105), CAST/EiJ (JAX 000928), PWK/PhJ (JAX 003715), and WSB/EiJ (JAX 001145).

Global (whole-body) knock-out mouse lines of *Arid1b* or *Chd8* were generated by first crossing conditional (floxed) mice from JAX—*Arid1b* (JAX 032061) (*38*) or *Chd8* (JAX 031555)—with female E2a-Cre mice (JAX 003724), and then back-crossing their progeny with wild-type mice (C57BL/6J; JAX 000664) to remove Cre. Genotypes of the final animals were rigorously validated by examining sequencing reads in our bulk Dip-C data—a form of whole-genome sequencing—and confirming heterozygous deletions of the floxed exons (i.e., read depths reduced to ~50% at the deleted exons, and reads spanning the deletion junctions).

10x Genomics Multiome

Cell nuclei were isolated from fresh frozen, postmortem human cerebellum or fresh mouse cerebellum following 10x Genomics Demonstrated Protocol "Nuclei Isolation from Complex Tissues for Single Cell Multiome ATAC + Gene Expression Sequencing" (CG000375 Rev B). Lysis time was optimized to 15 min (rather than 5 min) to prevent cell clumping at a later step (i.e., resuspension in PBS + 1% BSA + 1 U/uL RNase inhibitor for the second time).

For each donor/sample, ~10 k nuclei were loaded into a 10x Genomics Chromium Next GEM Single Cell Multiome ATAC + Gene Expression Kit following its user guide (CG000338 Rev E) with minor modifications: We used 10 and 8 PCR cycles (rather than 6 for ≥6 k nuclei) for cDNA amplification of human and mouse, respectively, and 20 uL (rather than 10 uL) of cDNA for GEX library preparation in Step 7.1 (**Table S2**).

Dip-C

Dip-C (*2*) was performed as in our previous study of the mouse forebrain (*5*), except that the 3C/Hi-C step was performed with the Arima 3C kit (which used a cocktail of 2 restriction enzymes for chromatin digestion). Detailed procedure can be found in our recent protocol paper (*4*). For some human experiments, protease inhibitor and RNase inhibitor were added during the nuclei isolation step to better preserve genome structure (**Table S2**); however, they had little effect on scA/B analysis (**Fig. S9**).

Pop-C

For each Pop-C reaction, a pool of cerebellar samples from 3–13 human donors or 2–8 mice were dropped into a single Dounce homogenizer (2–50 mL in size) and homogenized together (**Table S2**). We then proceeded with regular Dip-C (see section above). The only exception was the vDip-C sample, which was pooled after flow-sorting for mGreenLantern+ cells (i.e., 2 samples were sorted into the same FACS collection tube) and before the Arima 3C reaction.

vDip-C

To construct the vDip-C vector, mGreenLantern (*13*) and KASH (*29*) sequences were synthesized directly; the *Pcp2* promoter was obtained by PCR with previously published primers (*33*) from genomic DNA of C57BL/6J mice, and therefore differed by a few strain-specific nucleotides from the original version. A full plasmid map can be found in **Fig. S8**.

AAV was produced in the PHP.eB serotype. We injected $10^{11}$ vg (~10 uL) retro-orbitally into each young adult (P30–39) mouse (**Table S2**). For 3D genome modeling, we used F1 hybrids of MOLF/EiJ (JAX 000928; father) and C57BL/6J (JAX 000664; mother) because CAST/EiJ (JAX 000928) or its progeny does not allow PHP.eB to cross the BBB (*31*). Mice were sacrificed at least 2 weeks later (at P56, or 17–26 days later) to allow sufficient viral expression.

Cell nuclei were isolated as in regular Dip-C (*4*). Nuclei were diluted (to avoid high FACS event rates) in ~10 mL PBS + 1% BSA (+ 300 nM DAPI for one of the 2 samples) and flow-sorted on a BD FACSAria to isolate the 0.2–0.5% mGreenLantern+ cells (i.e., Purkinje cells) (**Fig. S8**). FACS data was plotted with FlowCal (version 1.3.0) (*44*). A total of ~70 k cells were obtained from the 4 mice. Sorted cells were processed with the Arima 3C kit (and flow-sorted again) as in regular Dip-C.

Bulk Dip-C

For bulk Dip-C of *Arid1b* and *Chd8* mutant and control mice, half of the Arima 3C product was used for bulk DNA extraction, following the same step as extracting DNA from digestion and ligation controls in our recent protocol paper (*4*). Sequencing libraries were prepared with Illumina DNA Prep (formerly known as Nextera DNA Flex), following its user guide (1000000025416 v10). We used a DNA input of 200 ng and the suggested 5 PCR cycles (**Table S2**).

Sequencing

Libraries were sequenced (paired-end 150 bp) on an Illumina NovaSeq (or in some cases, HiSeq) by Novogene. Multi-ome libraries were trimmed to read lengths specified by 10x Genomics.

Analysis of Multi-ome Data

*Pre-processing with Cell Ranger ARC and Seurat/Signac*

FASTQ files were pre-processed with Cell Ranger ARC (version 2.0.0) using “cell-ranger-arc count” and reference genomes “GRCh38-2020-A-2.0.0” and “mm10-2020-A-2.0.0” for human and mouse, respectively.

For each sample/donor, output files of Cell Ranger ARC were further pre-processed with Seurat (version 4) (*45*) and Signac (version 1.9.0) (*46*) following the tutorial “Joint RNA and ATAC analysis: 10x multiomic” (version: December 07, 2022; https://stuartlab.org/signac/articles/pbmc_multiomic.html). In particular, in R (version 4.1.2), the RNA filtered feature barcode matrix (“filtered_feature_bc_matrix.h5”) and ATAC fragments (“atac_fragments.tsv.gz”) were loaded with “CreateSeuratObject” and “CreateChromatinAssay.” Only cells with >500 and <25 k UMIs, >1 k and <100 k ATAC fragments, TSS enrichment score >1, and nucleosome signal <2 were kept—all were default parameters from the tutorial except >500 UMIs (rather than >1 k UMIs), which was the value used by (*18*). We used gene annotations “EnsDb.Hsapiens.v86”/ “” and “BSgenome.Mmusculus.UCSC.mm10” / “EnsDb.Mmusculus.v79” for human and mouse, respectively. RNA data was normalized with “SCTransform.”

*Cross-sample Integration and Analysis of the Transcriptome Portion with LIGER*

The transcriptome portion of the 7 human donors were integrated with LIGER (version 1.0.0) (*21*) following the tutorial “Joint definition of cell types from multiple scRNA-seq datasets” (version: March 31, 2020; http://htmlpreview.github.io/?https://github.com/welch-lab/liger/blob/master/vignettes/Integrating_multi_scRNA_data.html) with a minor modification: Direct input from Cell Ranger ARC did not work in our hands; instead, we first merged the 7 Seurat objects with “merge,” which merged raw RNA count matrices, and then converted the merged Seurat object into a LIGER object with “seuratToLiger” (with parameter “combined.seurat = TRUE”), which re-normalized the raw data by default.

In particular, the LIGER object was normalized and scaled with “normalize,” “selectGenes,” and “scaleNotCenter.” We then performed integrative non-negative matrix factorization (iNMF) with “optimizeALS” and parameter “k = 100”—the value used by (*18*); results with alternative values of “k” were shown in **Fig. S1**. Transcriptional cell types were identified with “quantile_norm,” “louvainCluster” (with parameter “resolution = 3”), and manual merging of the resulting clusters (“recode”). **Fig. 1B** were generated with “runUMAP” (with parameters “distance = 'cosine', n_neighbors = 30, min_dist = 0.3”—default parameters from the tutorial) and “plotByDatasetAndCluster.”

Cell type–specific marker genes were identified with “runWilcoxon” (with parameter “compare.method = "clusters"”). Genes shown in **Fig. 1C** were chosen based on test statistics, expression in other clusters, and whether they were known markers.

Correlated gene modules in immature granule cells (**Fig. 2A**) were identified by visually examining the heatmap produced by “plotClusterFactors” and finding factors that were highly active at the 4 immature granule cell stages. For each factor, genes were sorted by LIGER weights (“liger_object@W”).

*Joint Transcriptome and Chromatin Accessibility Analysis with ArchR*

For each sample/donor, RNA and ATAC data were jointly analyzed with ArchR (version 1.0.2) (*22*) following the tutorial "ArchR: 10x Multiome PBMCs" (https://greenleaflab.github.io/ArchR_2020/Ex-Analyze-Multiome.html). In particular, for maximum versatility, Arrow files were first generated from the ATAC output of Cell Ranger ARC ("atac_fragments.tsv.gz") using "createArrowFiles" without any initial filtering (Chapter 1.6 "Creating Arrow Files" of the manual). The RNA output of Cell Ranger ARC ("filtered_feature_bc_matrix.h5") was then added. Only cells with >500 UMIs (rather than 0; 500 is the value used by (*18*)), >2.5 k ATAC fragments (default in the tutorial), and TSS enrichment score >2 (rather than 6) were kept. We used reference genomes "hg38" and "mm10" for human and mouse, respectively. Doublet detection could not be performed because of a predominance of granule cells. We created an individual ArchR project for each sample/donor for subsequent analysis.

Iterative latent semantic indexing (LSI) was first performed separately on RNA ("GeneExpressionMatrix") and ATAC ("TileMatrix") with "addIterativeLSI" (with default parameters in the tutorial: "resolution = 0.2, sampleCells = 10000, n.start = 10", and additionally "varFeatures = 2500" for RNA). The RNA and ATAC dimensions were then combined with "addCombinedDims." Separate and joint UMAPs were created with "addCombinedDims" (with default parameter in the tutorial: "minDist = 0.8") and plotted with "plotEmbedding" in **Fig. 2B**.

Multi-omic cell types were identified with "addClusters" with resolutions 0.4, 0.4, 2, 1.5, 1, 0.4, and 1.5 (rather than the default 0.4 in the tutorial) for donors 1, 2, 3, 4, 5, 6, and 13, respectively, followed by manual merging of clusters.

Cell type–specific marker genes were identified with "getMarkerFeatures" from the RNA data ("GeneExpressionMatrix") following Chapter 7.3 "Identifying Marker Genes" of the manual.

To color LIGER-annotated transcriptional cell types on ArchR plots (**Fig. 2B**), we manually stored the correspondence between cell IDs with LIGER cluster IDs, added this information to each ArchR object as "archr_project$liger_clusters," and plotted with "plotEmbedding" (with parameter "name = liger_clusters").

ATAC peaks were called with "addReproduciblePeakSet" following Chapter 10.2 "Calling Peaks w/ Macs2" of the manual. The ATAC count matrix ("PeakMatrix") were generated with "addPeakMatrix" following Chapter 10.4 "Add Peak Matrix" of the manul.

To plot the accessibility of an individual ATAC peak on the ArchR plot (**Fig. 2C**), we manually created names for ATAC peaks by concatenating their chromosomes ("seqnames") and coordinates ("ranges"), and assigning these names to the "peakSet" (i.e., "archr_project@peakSet$name <- paste0(seqnames(archr_project@peakSet), "_", ranges(archr_project@peakSet))"). We then re-created the "PeakMatrix" with these new names using "addPeakMatrix" (with parameter "force = TRUE"), and plotted with "plotEmbedding" (with parameters such as "name = "chr16_54930458-54930958", imputeWeights = NULL").

Enrichment of TFBS motifs (**Fig. 2C**) was analyzed following Chapter 13.1 "Motif Deviations" of the manual.

Pseudotime trajectory of granule cell maturation (**Fig. 2C**) was analyzed following Chapter 16 "Trajectory Analysis with ArchR" of the manual. In particular, for each sample/donor, a trajectory was created from the joint RNA + ATAC UMAP embedding with "addTrajectory" (with parameter "spar = 2" for a smooth trajectory). Dynamic genes, peaks, and motifs were identified with "getTrajectory" (using "GeneExpressionMatrix," "PeakMatrix," and "MotifMatrix," respectively) and "plotTrajectoryHeatmap."

Published Data

*Single-cell Transcriptome of the Developing Human Cerebellum*

We downloaded raw data (FASTQ files) of the postnatal human portion (aged 0–42) of the preprint (*18*) from https://heidata.uni-heidelberg.de/dataset.xhtml?persistentId=doi:10.11588/data/QDOC4E.

We re-analyzed samples SN035, SN060, SN061, SN062, SN080, SN122, SN136, SN149, SN213, SN214, SN234, and SN283; we were unable to re-analyze SN233 and SN308 because of corrupt raw data and error in running Cell Ranger, respectively. In particular, FASTQ files were pre-processed with Cell Ranger (version 7.0.0). RNA count matrices were loaded into Seurat (version 4) using "CreateSeuratObject" without any additional filtering of cells. Data was normalized with "FindVariableFeatures," scaled with "ScaleData," and visualized with PCA and UMAP with the first 10 PCs (with paramter "dim = 1:10") (**Fig. 2D**).

Cell type–specific marker genes that were conserved between human, mouse, and opposum were from Supplementary Table 5 of (*18*). Pseudobulk expression was accessed from https://apps.kaessmannlab.org/sc-cerebellum-transcriptome/.

*Single-cell Transcriptome of the Adult Mouse Cerebellum*

Processed data from (*20*) were downloaded from https://singlecell.broadinstitute.org/single_cell/study/SCP795/.

Analysis of Dip-C Data

Dip-C data were analyzed as in our previous study (*5*), with a minor modification (cells with <50 k contacts were filtered out as empty wells, and >2 m contacts as cell aggregates; previously: cells with <20 k contacts as empty wells, no filters for cell aggregates) and the following additions:

*Visualization of Contact Maps*

Contact maps were visualized with Juicebox (*47*). Changes in contact maps (i.e., differential) were visualized by loading 2 maps ("A" and "B"), selecting the "A/B" option, and clicking "+" once (i.e., setting the scale to "20").

*Demultiplexing of Pop-C Data*

Sexes were demultiplexed by calculating the ratio between reads mapping to the X chromosome and reads mapping to autosomes, normalized by X chromosome and total autosome lengths ("X:A") with "samtools idxstats"—which exhibited a strong bimodal distribution and were separated by setting a threshold between the 2 modes (high X:A = female, low X:A = male). This could be alternatively achieved by calculating X:A for contacts rather than reads.

Mouse strains were demultiplexed by examining sequencing read pile-ups at strain-specific SNPs. Known SNPs were downloaded from the Sanger Institute Mouse Genomes Project FTP site (https://www.sanger.ac.uk/data/mouse-genomes-project/) as "[strain].mgp.v5.snps.dbSNP142.vcf.gz" files. Among the 8 founder stains of JAX DO, the 7 non-reference strains A/J, 129S1/SvImJ, NOD/ShiLtJ, NZO/HlLtJ, CAST/EiJ, PWK/PhJ, and WSB/EiJ differed from the mouse genome reference strain C57BL/6J by 4.9, 5.2, 5.1, 5.3, 20.7, 20.3, and 7.1 million SNPs, respectively.

Human donors were demultiplexed with souporcell (*24*) (version 2.4)—which we adapted from transcriptome data to 3D genome (or whole-genome sequencing in general) data—following the manual with minor modifications. In particular, we did not re-map the BAM files, but rather directly used our BWA MEM–mapped BAM files from Dip-C, where cell IDs were already recorded as the read group ("RG") BAM tag. PCR duplicates were removed from each BAM file with "sambamba markdup" (version 0.7.0). For each Pop-C experiment (note that we computationally pooled human experiments 2 (cerebellum) and 3 (cerebral cortex) because they contained nearly the same set of donors, and human experiments 8 (cerebellum) and 9 (cerebral cortex)) (**Table S2**), BAM files of all cells merged with "sambamba merge." Read count matrix at common SNPs in human populations ("common_variants_hg19.vcf.gz") was calculated with "vartrix" (version 1.1.20) (with parameter "--bam-tag RG" in addition to the default parameters in the manual "--mapq 30 --scoring-method coverage"). Finally, donors were demultiplexed with "souporcell" (with parameter "k" set to the number of pooled donors) and "troublet." We removed 2 cells ("dipc-human-experiment08-cerebellum_0843" and "dipc-human-experiment09-cortex_0426") that were classified at doublets after this step.

When demultiplexing the 13 donors from human experiments 8 and 9, we first demultiplexed by sex (see above) and then ran souporcell separately on the 2 sexes (4 female donors and 9 male donors), because running souporcell directly did not demultiplex 2 of the 13 donors.

In general, inclusion of empty wells (<50 k contacts) and cell aggregates (>2 m contacts) had no impact on souporcell except in one case, where a cell ("dipc-human-experiment06-cerebellum_0059") was classified as a singlet only when empty wells and cell aggregates were filtered out before running souporcell.

*scA/B Dynamics across Maturation Stages*

To identify dynamic genomic regions during granule cell maturation, we calculated the mean scA/B of each 1-Mb autosomal regions at each of the 5 maturation stages, and selected the top 10% regions with the highest between-stage variance in MATLAB (version R2022b). We then performed hierarchical clustering of these dynamic regions based on their mean scA/B at the 5 maturation stages, with "linkage_data = linkage(scab_matrix(dynamic_regions_ids, :), 'average', 'correlation')." To plot their centered scA/B heatmap (**Fig. 5A**), we ordered dynamic regions based on their hierarchical clustering, with "outperm" from "[H, ~, outperm] = dendrogram(linkage_data, 0)."

To analyze scA/B of individual genes (**Fig. 5B**), mid-points of genes were calculated from the GENCODE annotations “gencode.v42lift37.annotation.gtf.gz” and “gencode.vM25.annotation.gtf.gz” for human and mouse, respectively.

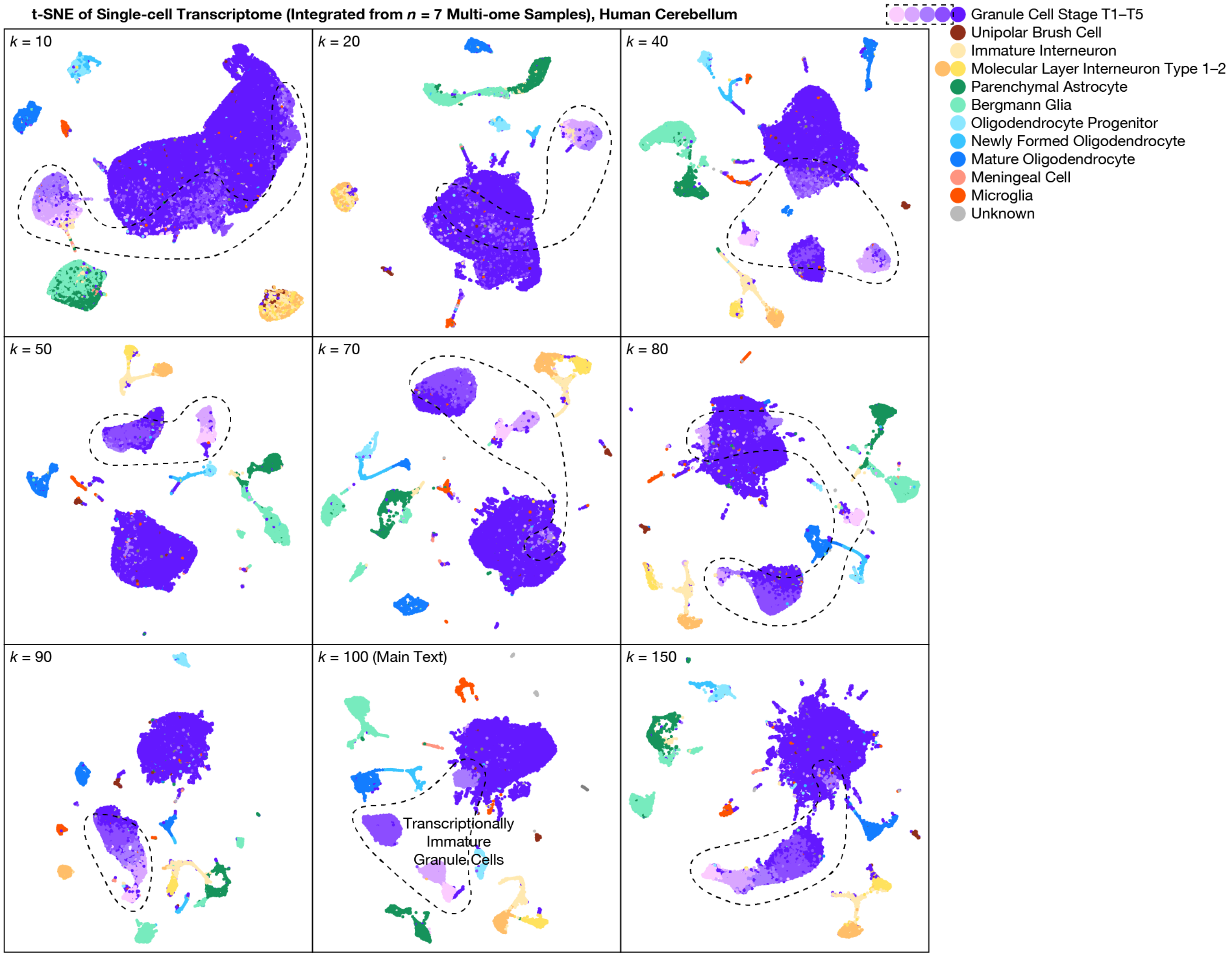


**Fig. S1. Integrative transcriptome analysis of our human multi-ome samples with alternative parameter values.** Similar to **Fig. 1B** but with alternative values of the LIGER parameter $k$ (i.e., the number of LIGER factors): $k = 10, 20, 40, 50, 70, 80, 90, 100$ (same as the main text), and 150. Cells were colored by transcriptional cell types from the main text (i.e., identified with $k = 100$). Regardless of the value of $k$, transcriptionally immature cerebellar granule cells (stage T1–T4: dashed outlines; lighter shades of purple) were consistently visualized as relatively distinct clusters on the t-SNE plot, although boundaries were variable because of the continuous nature of their maturation.

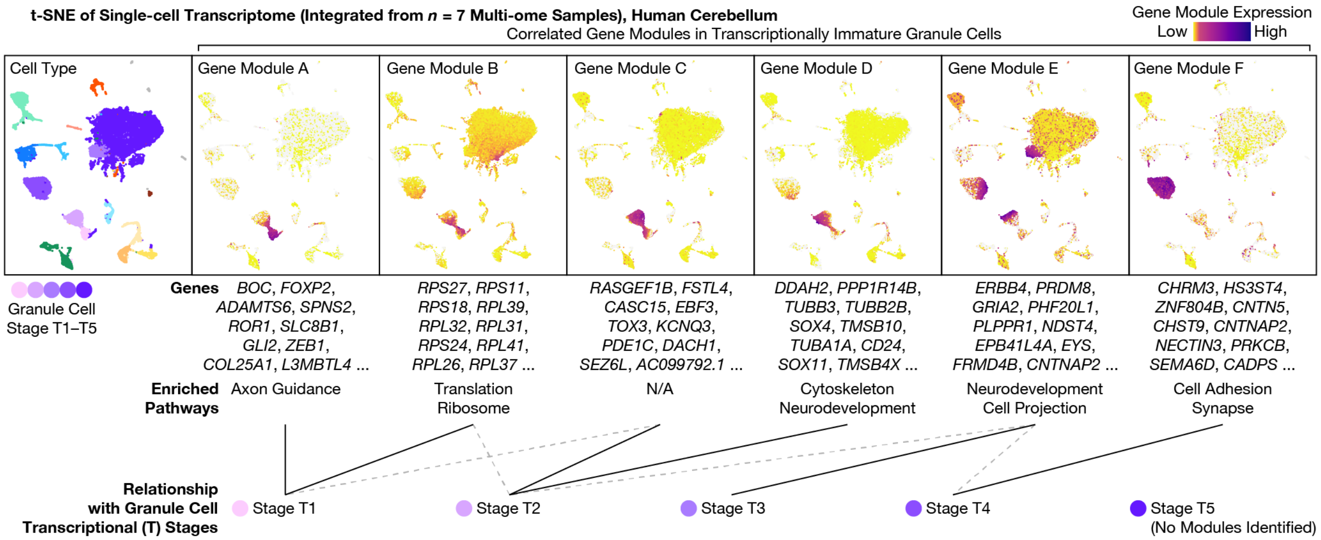


**Fig. S2. Correlated gene modules in transcriptionally immature cerebellar granule cells.** We identified 6 correlated gene modules A–F from LIGER factors that were specifically expressed at each transcriptional (T) stage of granule cell maturation. The top row shows the expression pattern of each module, quantified by LIGER metagene expression (weighted average of genes in each module) on the t-SNE plot. The middle rows show the top 10 genes (ranked by LIGER weights) and enriched pathways (summarized GO terms for the top 50 genes) for each module. The bottom row connects each module to its corresponding T stages (solid line: most expressed; dashed line: also expressed). Module A was enriched for the pathway of neuron projection guidance (FDR = $3 \times 10^{-4}$), included genes such as *BOC*, *FOXP2*, *ADAMTS6*, and was expressed at stage T1. Modules B (various ribosome subunits) and C were expressed at both stages T1–T2. Module D was enriched for cytoskeleton (FDR = $2 \times 10^{-9}$) and nervous system development (FDR = $9 \times 10^{-8}$), included genes such as *TUBB3/2B/1A*, *SOX4/11* (transcription factors implicated in Coffin-Siris syndrome), *TSMB10/4X*, and was expressed at stage T2. Module E was enriched for telencephalon development (FDR = $7 \times 10^{-5}$) and cell projections (FDR = $3 \times 10^{-4}$), included genes such as *ERBB4*, *PRDM8* (a histone H3K9 methyltransferase implicated in epilepsy), *GRIA2*, and was expressed at stage T3. Finally, module F was enriched for cell adhesion (FDR = $8 \times 10^{-4}$) and synapses (FDR = $4 \times 10^{-7}$), included genes such as *CHRM3* (a muscarinic acetylcholine receptor), *HS3ST4*, *CNTN5* (a contactin implicated in Coffin-Siris), *CNTNAP2* (a neurexin implicated in multiple neurodevelopmental disorders including autism), and was expressed at stage T4—the most abundant immature type. Note that LIGER did not identify gene modules for the mature stage T5.

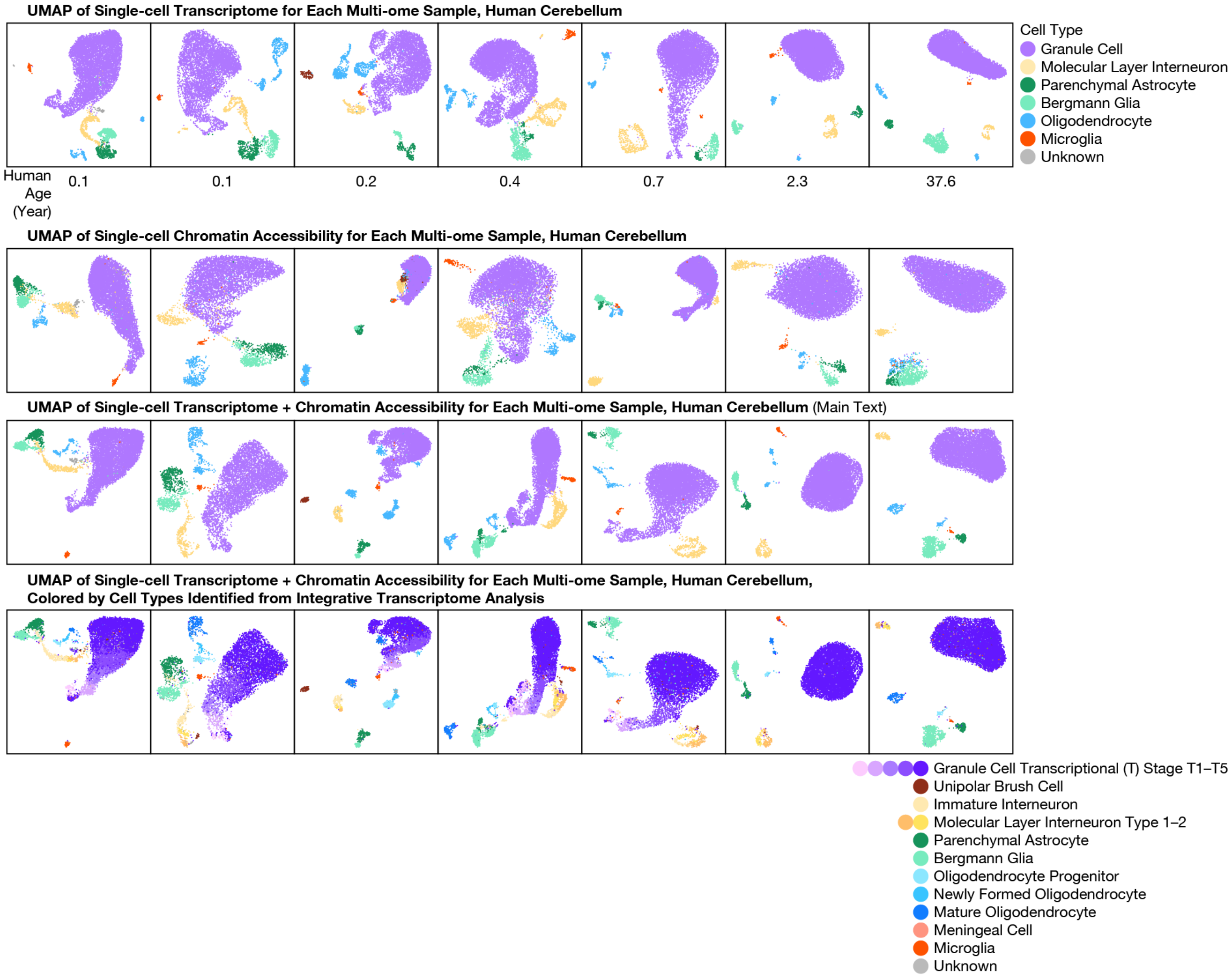


**Fig. S3. Alternative visualizations of each donor in our human multi-ome atlas.** Similar to **Fig. 2B** but visualized with the transcriptome portion only (first row), the chromatin accessibility portion only (second row), transcriptome and chromatin accessibility jointly (third row; same as the main text), or transcriptome and chromatin accessibility jointly but colored by transcriptional cell types from integrative transcriptome analysis (i.e., by LIGER, in **Fig. 1B**) (last row).

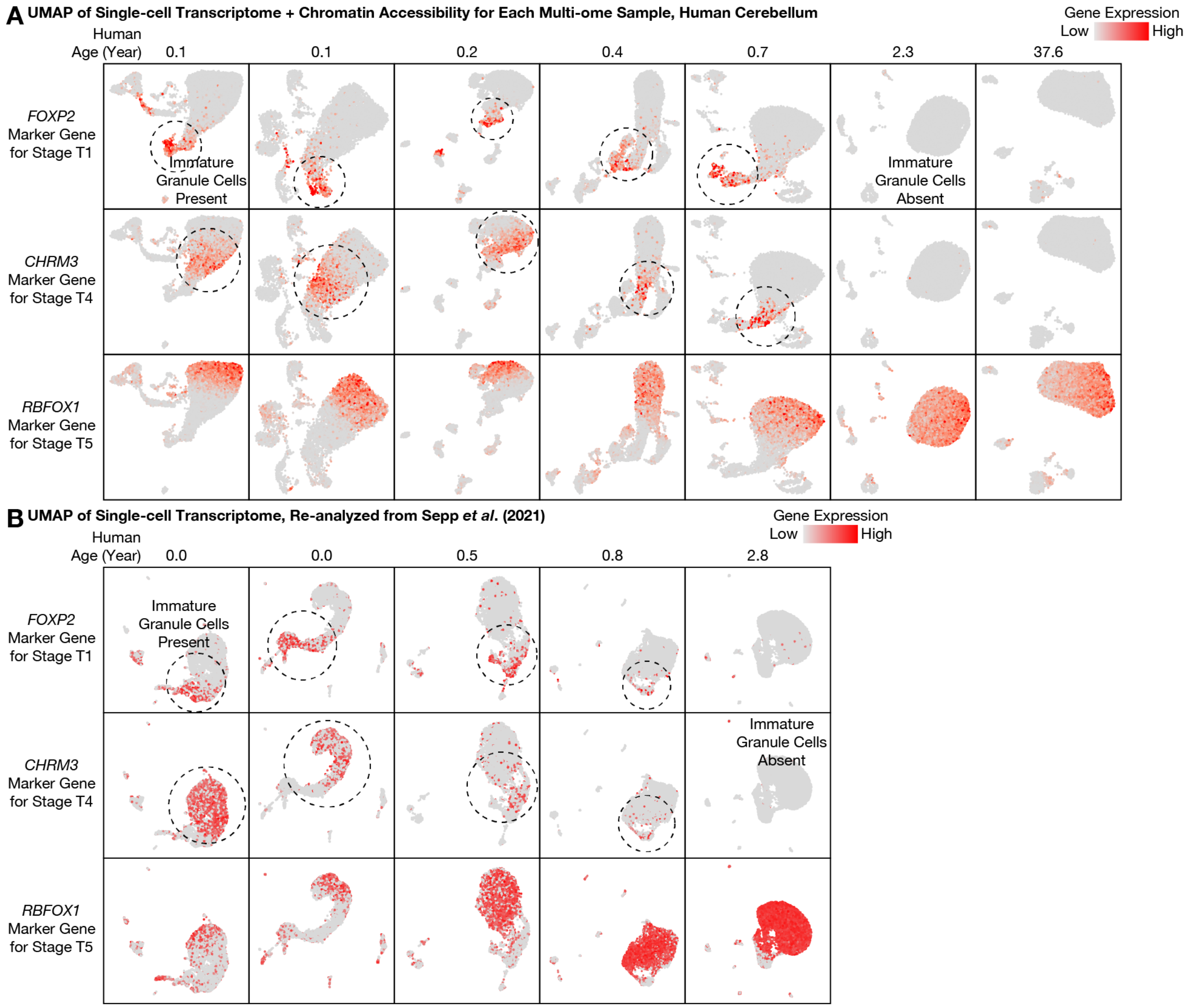


**Fig. S4. Expression patterns of transcriptionally immature and mature granule cell marker genes in each donor of our human multi-ome atlas and in published transcriptome data. (A)** Similar to **Fig. 2B** but for all 7 donors. **(B)** Similar to **(A)** but for data re-analyzed from a single-cell transcriptome atlas preprinted while this paper was in preparation (*18*).

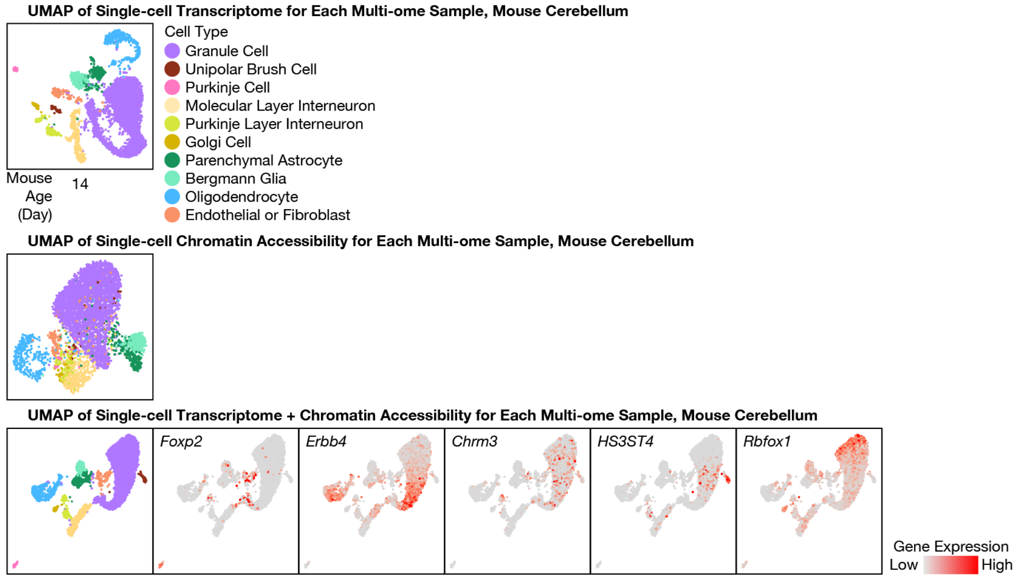


**Fig. S5. A continuum of granule cells with maturing transcriptome and chromatin accessibility in the newborn mouse cerebellum.** Similar to **Fig. S3** and **Fig. S4A** but for our mouse multi-ome sample (P14).

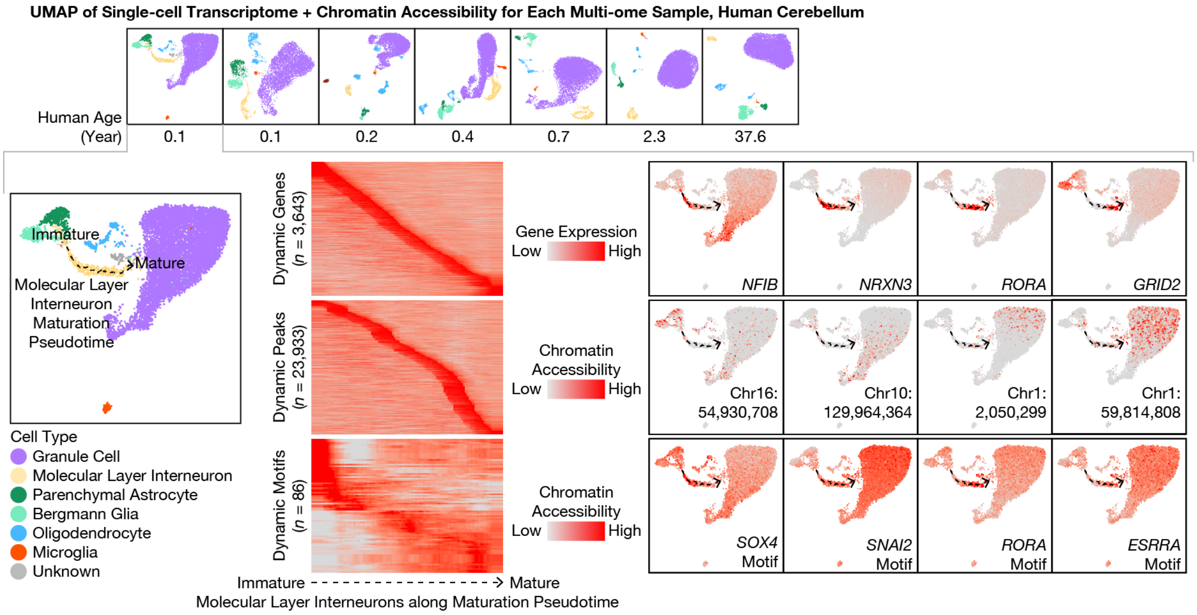


**Fig. S6. Maturation pseudotime analysis of molecular layer interneurons (MLIs) in our human multi-ome atlas.** Similar to **Fig. 2B** but for MLIs.

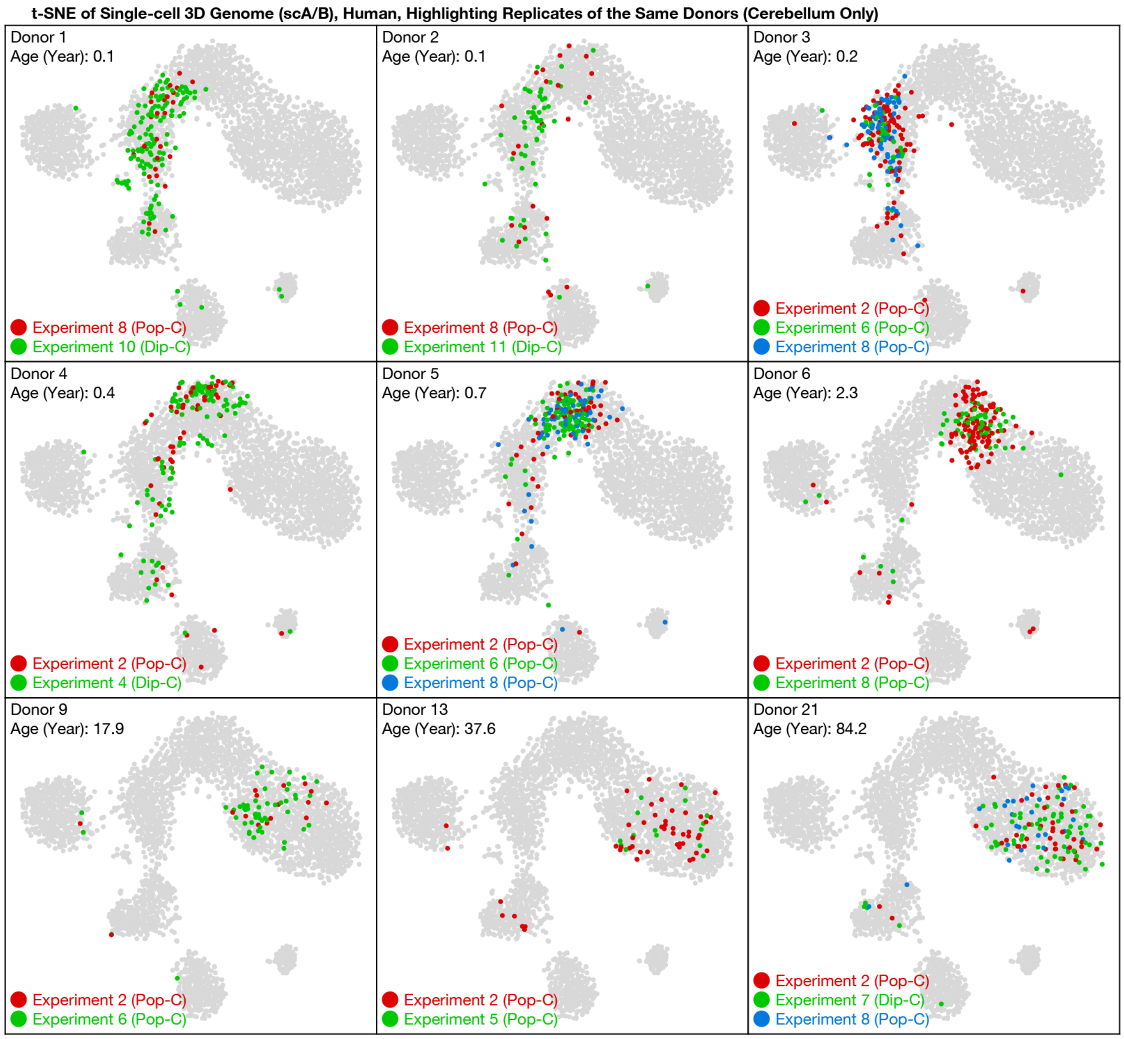


**Fig. S7. Reproducible 3D genome profiling with Pop-C and Dip-C between replicates.** Similar to **Fig. 3A** but highlighting different replicates (red, green, and blue) of the same donors on the t-SNE plot.

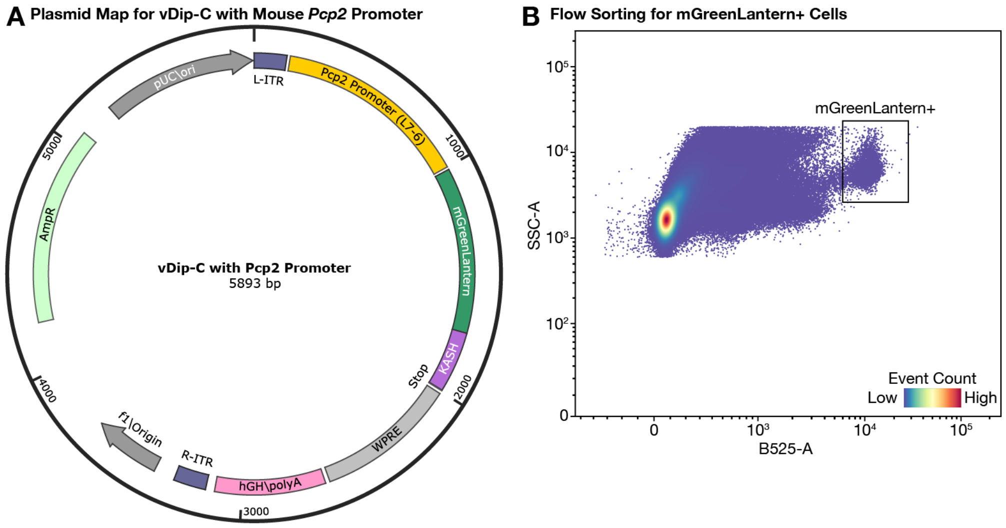


**Fig. S8. Plasmid map and flow sorting for vDip-C to isolate rare Purkinje cells. (A)** Plasmid map for the vDip-C vector. The promoter of the mouse *Pcp2* gene (844 bp) was taken from (*33*) ("L7-6"). The coding sequence of mGreenLantern (717 bp) was taken from (*28*). The KASH nuclear membrane–location tag (264 bp) was taken from (*29*). L/R-ITR: left/right inverted terminal repeats. WPRE: woodchuck hepatitis virus post-transcriptional regulatory element. AmpR: ampicillin resistance. **(B)** Representative FACS plot to isolate mGreenLantern+ Purkinje cells (boxed, yellow dots) from the adult mouse cerebellum. SSC-A: side scattering, area. B525-A: band-pass 525 nm fluorescence, area.

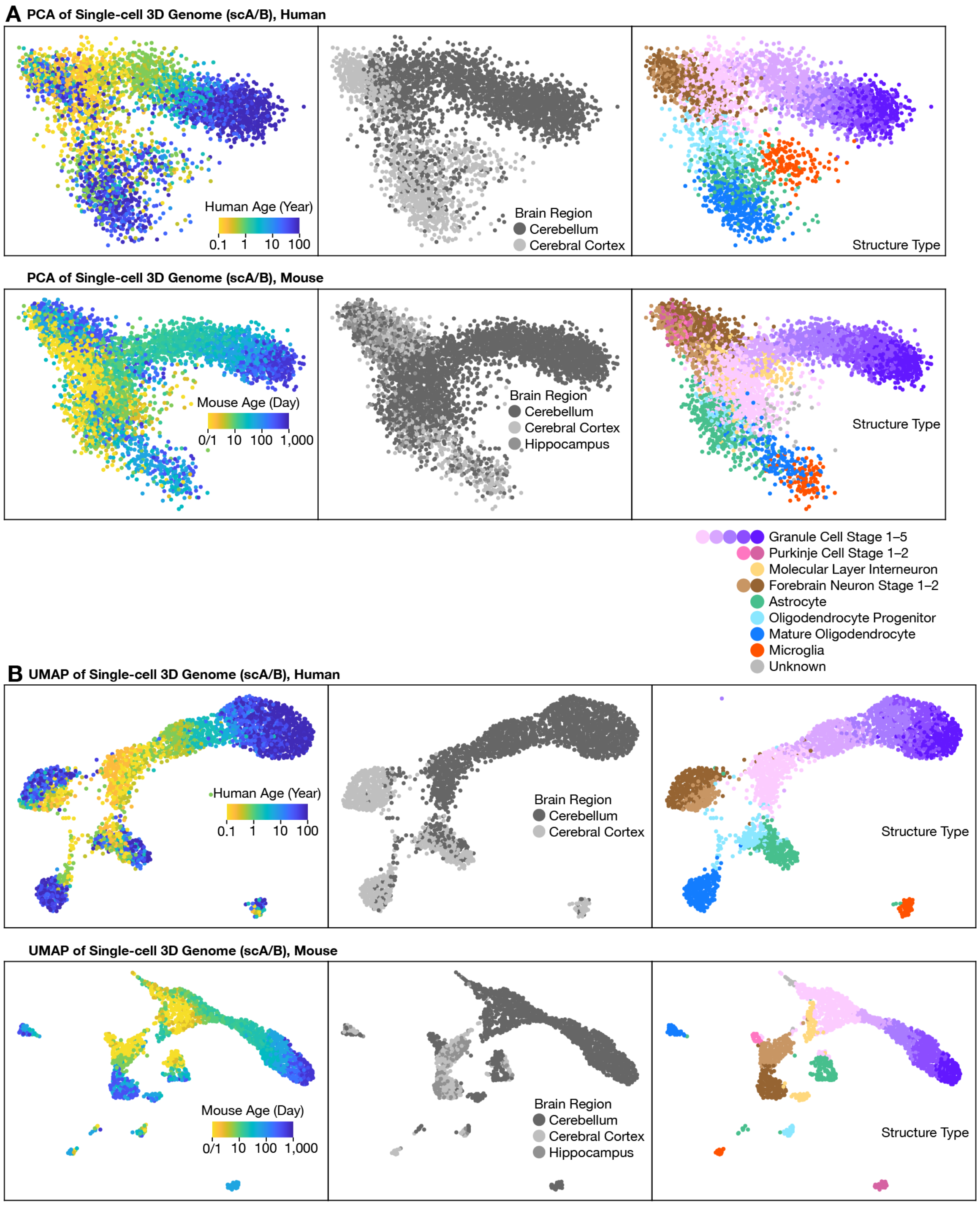


**Fig. S9. Alternative visualizations of our 3D genome atlas with PCA or UMAP of scA/B. (A)** Similar to **Fig. 3C** and **Fig. 3D**, but with PCA rather than t-SNE. **(B)** Similar to **Fig. 3C** and **Fig. 3D**, but with UMAP rather than t-SNE.

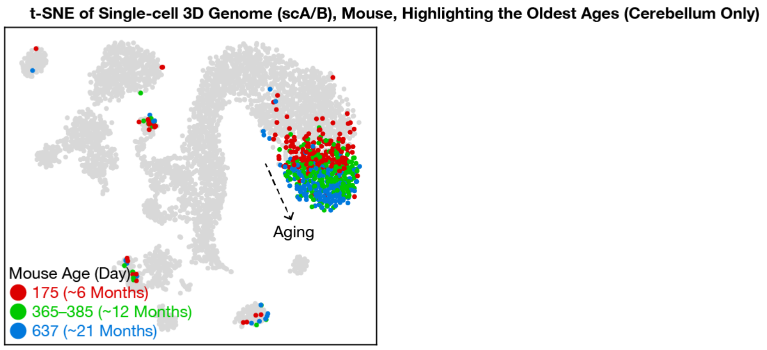


**Fig. S10. Continued 3D genome remodeling in the aging mouse cerebellum.** Similar to **Fig. 3D** but highlighting only the oldest ages: P175 (~6 months; red), P365–385 (~12 months; green), and P637 (~21 months; blue).

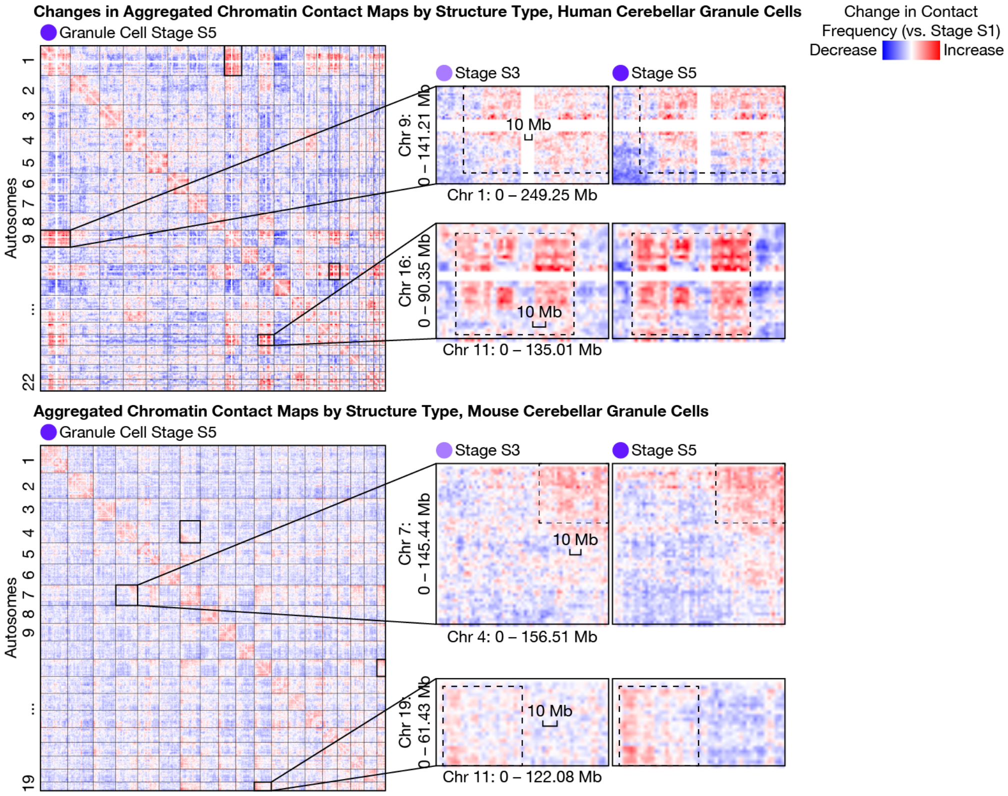


**Fig. S11. Differential inter-chromosomal chromatin contact maps of granule cells over the lifespan.** Similar to **Fig. 4C** but showing changes in contact maps (with respect to structural (S) stage S1). Bin size was 6 Mb (human genome-wide), 5 Mb (mouse genome-wide), or 3 Mb (zoom-in).

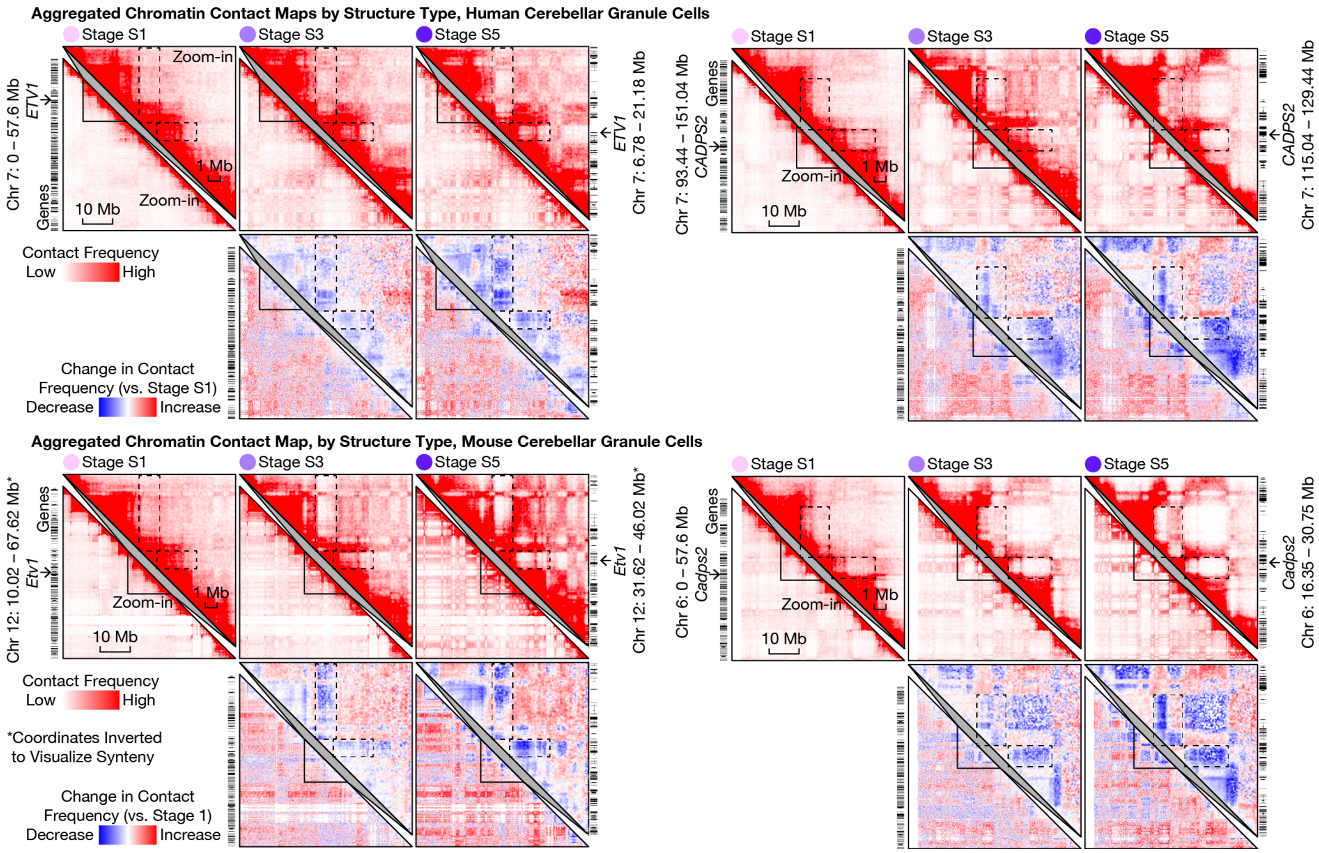


**Fig. S12. Life-long 3D genome rewiring of other mature granule cell–specific marker genes.** Similar to **Fig. 5C** but for 2 other marker genes, *ETV1* (a cancer-implicated transcription factor; left) and *CADPS2* (an autism-implicated calcium-dependent secretion activator; right).

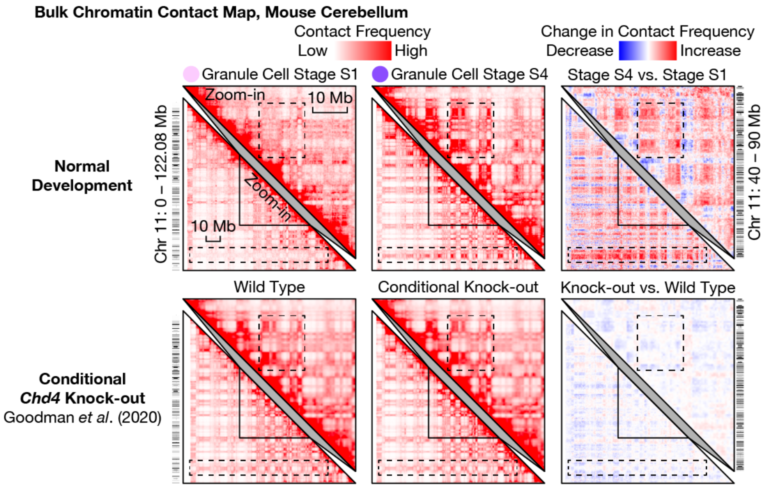


**Fig. S13. Life-long 3D genome changes of granule cells are resistant to conditional knock-out of *Chd4*.** Similar to **Fig. 5D** but with *Chd4* knock-out rather than *Arid1b* or *Chd8*. Bulk Hi-C data after conditional (driven by *Gabra6*-Cre), homozygous deletion of *Chd4* was from (*15*). Although *Chd4* deletion moderately changed 3D genome (bottom right), these changes had little overlap with and had a much smaller magnitude than our observed architectural maturation (top right). Bin sizes was 250 kb (top) or 500 kb (because published data did not contain 250 kb resolution; bottom).

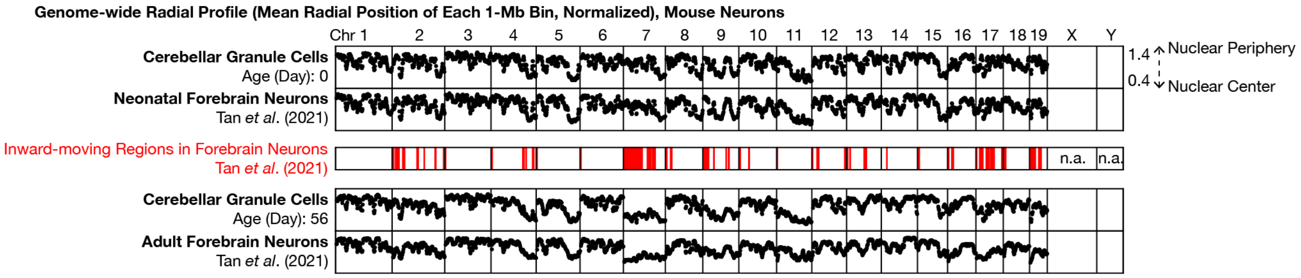


**Fig. S14. Inward movement of many genomic regions occurs in both forebrain neurons and cerebellar granule cells, despite the lack of non-CpG methylation in the latter.** Similar to Fig. 4B of our previous study in the mouse cortex and hippocampus (*5*), but plotting cerebellar granule cells (first row: neonatal; fourth row: adult) and forebrain neurons (second row: neonatal; fifth row: adult) side by side. Despite a lack of neuron-specific non-CpG methylation (*16*)—which we found to occur simultaneously with and anti-correlate with inward genome movement in forebrain neurons—cerebellar granule cells still exhibited inward genome movement (e.g., Chr 7).

**Table S1. Information about each human donor.**

**Table S2. Information about each multi-ome and 3D genome experiment.**

**Table S3. Number of cells from each human donor in each 3D genome experiment.**

**Table S4. Information about each Dip-C cell.**

**Table S5. Lists of cell type–specific marker genes.**
